# Defining the early stages of intestinal colonisation by whipworms

**DOI:** 10.1101/2020.08.21.261586

**Authors:** María A. Duque-Correa, David Goulding, Faye H. Rodgers, Claire Cormie, Kate Rawlinson, J. Andrew Gillis, Allison J. Bancroft, Hayley M. Bennett, Magda Lotkowska, Adam J. Reid, Anneliese Speak, Paul Scott, Nicholas Redshaw, Catherine McCarthy, Cordelia Brandt, Catherine Sharpe, Caroline Ridley, Judit Gali Moya, Claudia M. Carneiro, Tobias Starborg, Kelly S. Hayes, Nancy Holroyd, Mandy Sanders, David J. Thornton, Richard K. Grencis, Matthew Berriman

**Affiliations:** Wellcome Sanger Institute, Wellcome Genome Campus, Hinxton, CB10 1SA, UK; Department of Zoology, University of Cambridge, Cambridge, CB2 3EJ, UK; Lydia Becker Institute of Immunology and Inflammation, Wellcome Trust Centre for Cell Matrix Research and Faculty of Biology, Medicine and Health, University of Manchester, Manchester, M13 9PT, UK; Faculty of Biology, University of Barcelona, Barcelona, 08028, Spain; Immunopathology Laboratory, NUPEB, Federal University of Ouro Preto, Campus Universitario Morro do Cruzeiro, Ouro Preto, MG, 35400-000, Brazil

## Abstract

Whipworms are large metazoan parasites that inhabit distinct multi-intracellular epithelial burrows described as syncytial tunnels, in the large intestine of their hosts. How first-stage larvae invade host epithelia and establish infection remains unclear. Here, we investigate early infection events both using *Trichuris muris* infections of mice and murine caecaloids, the first *in-vitro* system for whipworm infection. We show that larvae degrade the mucus layers to access epithelial cells. In early syncytial tunnels, larvae are completely intracellular but woven through multiple live enterocytes and goblet cells. We also use single cell RNA sequencing for the first time to describe the mouse caecum. From infected caeca, the transcriptome data reveal the progression of infection results in cell damage and an expansion of enterocytes with a type-I interferon (IFN) signature, characterised by the expression of *Isg15*, instigating the host immune response to the whipworm and tissue repair. Our results unravel intestinal epithelium invasion by whipworms and reveal new specific interactions between the host and the parasite that allow the whipworm to establish its multi-intracellular niche.

## Introduction

Human whipworms *(Trichuris trichiura)* infect hundreds of millions of people and cause trichuriasis, a major Neglected Tropical Disease with high chronic morbidity and dire socio-economic consequences in affected countries^1, 2^. Although *T. trichiura* is experimentally intractable, a mouse model of infection with the natural rodent-infecting species *T. muris* closely mirrors infections in humans^3^, making this species distinctive as the only major soil transmitted helminth with a direct mouse counterpart.

Whipworms live in the caecum and proximal colon of their hosts and have a unique life cycle strategy where they establish a multi-intracellular niche within intestinal epithelial cells (IECs)^3–5^. In this niche, whipworms can remain for years causing chronic infections^1, 2^. Infection with whipworms follows ingestion of eggs from the external environment^1–3^. Upon arrival in the caecum and proximal colon, eggs hatch in a process mediated by the host microbiota^3, 6^ *(Fig. 1a)*. Within hours, motile first-stage (L1) larvae released from the eggs enter the intestinal epithelia (IE) at the bottom of the crypts of Lieberkühn^4, 7–9^ *(Fig. 1A)*. To reach this location, L1 larvae need to overcome barriers that are known to protect the crypt base, including the mucus layers covering the epithelium and the continuous stream of fluid that flushes from the crypt to the lumen^10^. To date, the physical and molecular cues directing the larvae to the crypt base and mediating their penetration through the overlying mucus into the IE are unknown.

**Figure 1.**
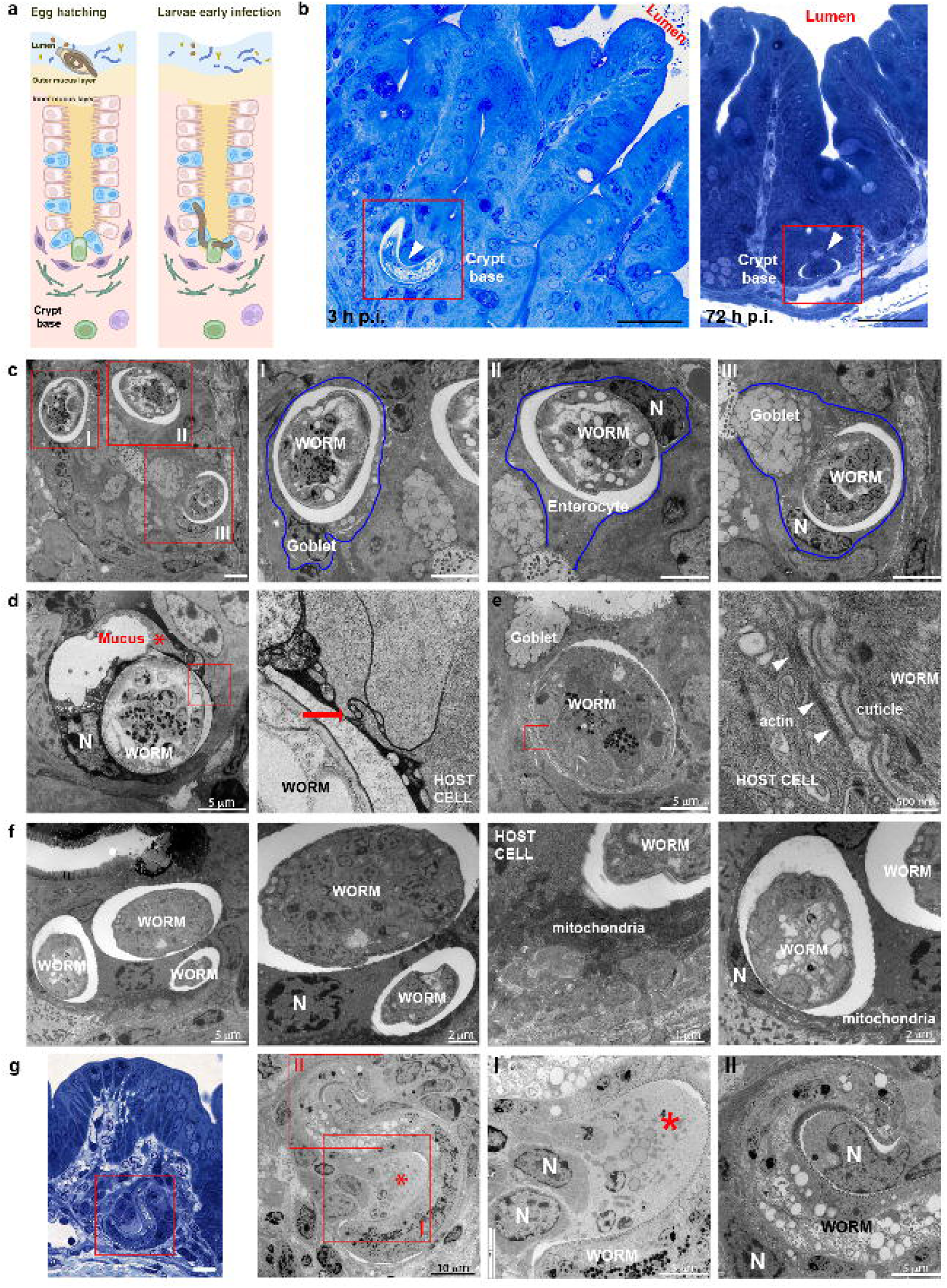
Whipworm L1 larvae infects enterocytes and goblet cells at the bottom of the crypts of Lieberkühn in the caecum. **a** Illustration of the processes of egg hatching at the caecal lumen and larvae infection of the IE at the base of the crypts. **b** Images of toluidine blue-stained transverse sections from caecum of mice infected with *T. muris* (3 and 72 h p.i.), showing whipworm larvae (arrowhead) infecting cells at the base of crypts. Scale bar 30μm. **c-g.** TEM images of transverse sections from caecum of mice infected with *T. muris.* **c** Whipworm L1 larvae infecting goblet cells (insets I and III) and enterocytes (inset II) at 3 h p.i. Scale bar 5μm. Blue lines show the cellular membranes of the host cells. **d-e** Larvae infecting goblet cells in the caecum of mice 3 h p.i., note: **d** potential mucus discharge (red asterisk) and tannic acid staining (black secretion) revealing complex carbohydrate in the host cell cytoplasm and between cells (inset, red arrow); **e** host cell cytoskeleton reorganization of actin filaments adjacent and parallel to the cuticle of the larvae (inset, white arrowheads). **f** Larvae infecting several IECs in the caecum of mice at 24 h p.i. Host cells display DNA condensation and fragmentation (pyknotic nuclei, characteristic for the onset of apoptosis) and numerous mitochondria. **g** Toluidine blue-stained (scale bar 20μm) and TEM images of transverse sections from caecum of mice infected with *T. muris*, showing a syncytial tunnel formed by L1 whipworm larvae through IECs (72 h p.i.) and depicting liquefaction of cells (inset I, red asterisk) and nuclei in early stages of apoptosis (inset II). N, nuclei.

To accommodate themselves completely inside the IE, the larvae burrow through several IECs creating multinucleated cytoplasmic masses that have been described as syncytial tunnels^11^. The syncytium is thought to provide a sheltered environment and a continuous source of nutrients for the worm, as it moults four times to reach adulthood^5, 11, 12^. Whilst the syncytial tunnels are only visible for the first time at the L3 larval stage, around day 21 post infection (p.i.), the biology and the mechanisms in which the multicellular epithelial burrows are generated are not understood^1^. Due to the lack of experimental accessibility during the first days of infection, the impact of the tunnels is controversial; it is not known whether IECs forming the tunnels are dead, or whether they are alive and interacting with the parasite to orchestrate the development of immune responses^5, 11, 13, 14^.

Here, we investigated the cellular and molecular processes mediating invasion of the IE by whipworm larvae and their establishment within multicellular epithelial tunnels during the first three days of infection. We used a combination of mouse infections, together with a new *in vitro* model comprising murine caecal oraganoids (caecaloids) infected with*T. muris* L1 larvae, to examine infection biology. We found that L1 larvae degrade the mucus layer and penetrate the underlying IE. We also observed that early syncytial tunnels are composed of enterocytes and goblet cells that are alive and actively interacting with the larvae during the first hours of the infection. Progression of infection results in damage to the host epithelium, which responds with an expression signature of type-I IFN signalling dominated by *Interferon-stimulated gene 15* (*Isg15*), an alarmin that initiates immune and tissue repair responses^15–17^. Collectively, our work unravels mechanisms involved in the invasion and colonisation of IE by whipworms through the formation of syncytial tunnels, and the early host IE-parasite interactions that can lead to the initiation of immune responses to the worm.

## Results

### Enterocytes and goblet cells at the bottom of the crypts of Lieberkühn are the host cells of whipworm L1 larvae

Light microscopy studies dating back 40-50 years have shown *T. muris* L1 larvae infecting cells at the base of the crypts of Lieberkühn in the first hours of an infection^4, 7, 8^. Confirming these findings, we found L1 larvae invading IECs in the crypt bases of the caecum and proximal colon of *T. muris*-infected mice as early as three hours p.i. *(Fig. 1b; Supplementary Fig. 1a-c).* Transmission electron microscopy (EM) revealed L1 larvae infecting enterocytes and goblet cells *(Fig. 1c)*. Larvae were in direct contact with the IEC cytoplasm as no cell membrane could be seen between the whipworm cuticle and the cytoplasm *(Figs. 1c-g)*. We found larvae displaced cellular organelles *(Figs. 1c-g)* and burrowed through mucin secretory granules of goblet cells, possibly causing mucus discharge *(Fig. 1d)*. Tannic acid staining revealed complex carbohydrate in the immediate vicinity surrounding the larvae, both between host cells and in bordering host cell cytoplasm, likely to be secretions from the worm or the result of disrupting goblet cells *(Fig. 1d, inset).* Infected cells reorganized their cytoskeleton around the worm *(Fig. 1e, inset).* With infection progression, at 24 and 72 h p.i., chromatin was visibly condensed and fragmenting indicating onset of apoptosis *(Figs. 1f* and *g)*, host cells show numerous mitochondria *(Figs. 1f)* and some host cells were liquified *(Figs. 1g, inset I).* Our observations were often limited to histological sections with a transverse view through a single slice of the worm within an IEC. Indeed, due to the intricate topography of the multicellular epithelial niche of the larvae, obtaining longitudinal sections of the complete worm inside its niche proved extremely challenging. However, serial block face SEM allowed us to capture the entire syncytial tunnel formed by L1 larvae *(Supplementary video 1 and 2)* and revealed that by 3 h p.i., larvae were completely intracellular. A typical syncytial tunnel was composed of approximately 40 IECs, with 75% of those cells being enterocytes and 15% goblet cells *(Supplementary Fig. 1d)*. Our results suggest that close interactions of enterocytes and goblet cells with L1 larvae are critical during invasion and colonisation of the IE by whipworms.

### Caecaloids provide an *in-vitro* infection model that reveals the intricate path of early syncytial tunnels

Although illuminating, serial block face SEM is technically demanding and is constrained by the need to find the small percentage of infected IECs (< 1%) in the total caecal epithelia and by the intracellular location of the L1 larvae at the bottom of crypts. Moreover, the lack of genetic tools to generate fluorescent larvae and of an *in vitro* culture system (cell lines do not support infection by whipworm L1 larvae) have severely hampered investigations on the early stages of intestinal colonisation by whipworms. Hence, to further examine the processes of invasion and formation of the syncytial tunnels, we developed the first *in vitro* whipworm infection model using caecaloids^18^. Caecaloids cultured in an open conformation using transwells *(Supplementary Fig. 2a)* generated self-organizing structures that resembled crypts present in the caecum. The structures comprised tight centres of proliferating stem and trans-amplifying (TA) cells, surrounded by differentiated absorptive, goblet, enteroendocrine and tuft cells, with a mucus layer overlaying the IECs; and polarised microvilli *(Supplementary Fig. 2b-k).* L1 larvae obtained by hatching *T. muris* eggs in the presence of *E. coli,* to simulate microbiota exposure^6^, were directly cultured with the IECs *(Supplementary Fig. 2a)*. We observed L1 larvae infecting IECs in caecaloids as evidenced by enterocyte microvilli staining above the worm *(Fig. 2a)* and images showing the larvae woven through multiple IECs *(Fig. 2b)*. We captured invasion of goblet cells by L1 larvae with SEM *(Fig. 2c),* and by TEM we found larvae within the cytoplasm of enterocytes and goblet cells of the caecaloids *(Fig. 2c).* Our caecaloid system also revealed the intricate path of the tunnels formed by L1 larvae burrowing through IECs *(Fig. 2d, Supplementary videos 3 and 4).* Together, these images show L1 larvae infecting IECs for the first time *in vitro,* effectively reproducing *in vivo* infection. The caecaloid model enabled the entirety of the L1 larva, its host cells, and the trail of the syncytial tunnels to be visualised.

**Figure 2.**
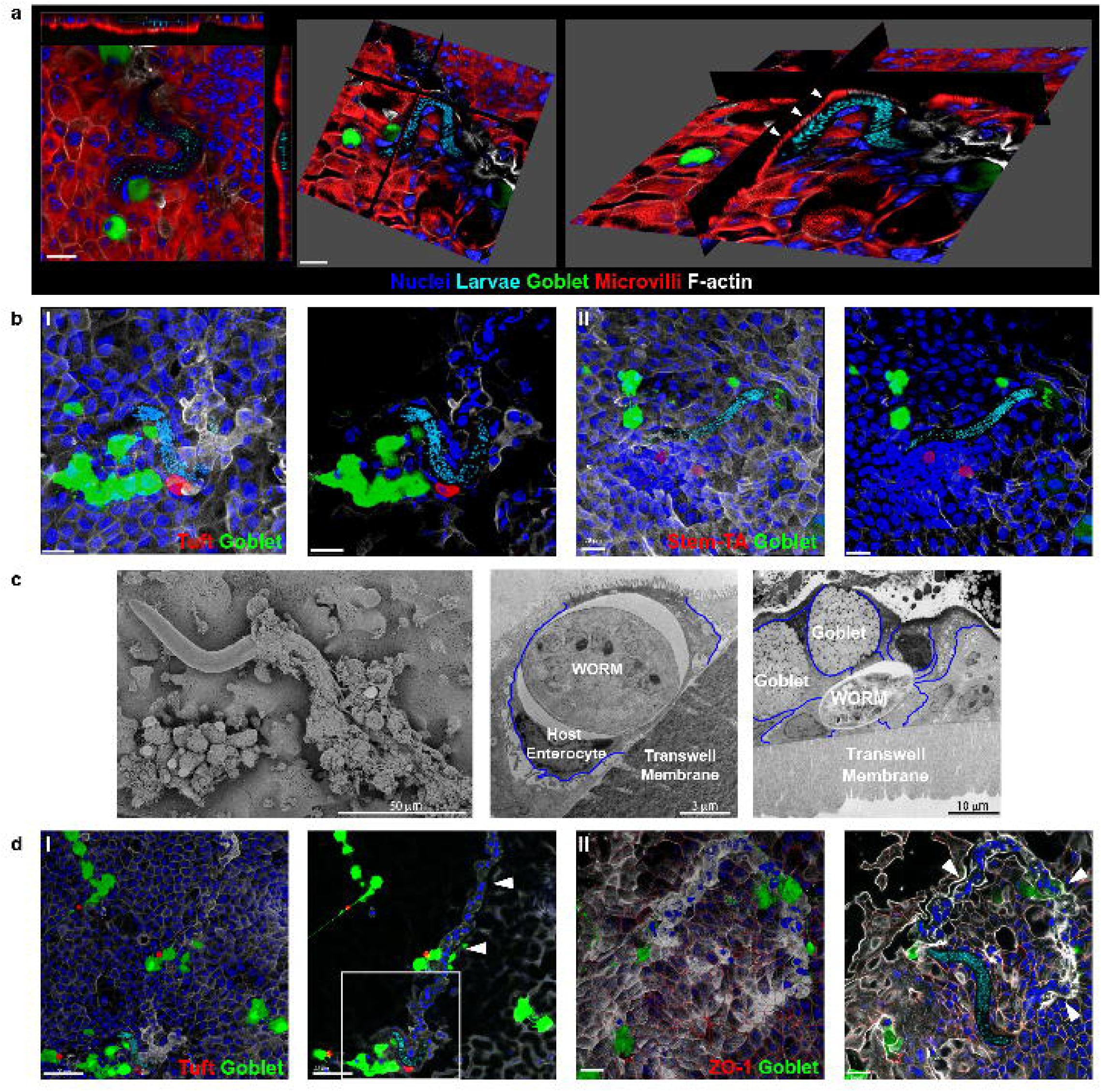
Caecaloid - *T. muris in vitro* model reproduces *in vivo* infection and reveals intricate path of syncytial tunnels burrowed by whipworm L1 larvae. **a-b** Confocal immunofluorescence (IF) images of caecaloids infected with whipworm L1 larvae for 24 h. **a** Orthogonal slice visualising enterocyte microvilli (villin staining in red) above the larvae (white arrowheads). Scale bar 20μm. **b** Complete z-stack projection and selected and cropped volume showing larvae infecting different IECs. In red, **(I)** Dclk-1, marker of tuft cells; **(II)** Ki-67, marker of proliferating cells, stem and TA cells. In green, the lectins *Ulex europaeus* agglutinin (UEA) and *Sambucus nigra (*SNA) bind mucins in goblet cells; in blue and aqua, DAPI stains nuclei of IECs and larvae, respectively; and in white, Phalloidin binds to F-actin. Scale bar 20μm. **c** Scanning and transmission EM images from caecaloids infected with *T. muris* for 24 h, showing whipworm L1 larvae within enterocytes and goblet cells. Blue lines show the cellular membranes of the host cells. **d** Complete z-stack projection and selected and cropped volume of confocal IF images of syncytial tunnels (white arrowheads) in caecaloids infected with L1 whipworm larvae for 24 h. In red, **(I)** Dclk-1, marker of tuft cells; **(II)** ZO-1 protein, binding tight junctions; in green, the lectins UEA and SNA bind mucins in goblet cells; in blue and aqua, DAPI stains nuclei of IECs and larvae, respectively; and in white, Phalloidin binds to F-actin. Scales bar for **(I)** 50μm, and **(II)** 20μm. Inset in **I** corresponds to figure **b (I)**.

### Whipworm larvae invade the caecal epithelium by degrading the overlaying mucus layer

After hatching and to counter host peristalsis, whipworm L1 larvae rapidly reach the bottom of the crypt and invade the IECs^8^. But first, the larvae must traverse the outer and inner mucus layers overlaying the IE. Despite their motility, mucus can aggregate around whipworm larvae, blocking their advance towards the IE. An additional mechanism is therefore required for larval traversal of the mucus layers. Many intestinal pathogens have evolved enzymes to degrade the mucin oligosaccharides via glycosidases, exposing the mucin peptide backbone to proteases^19^. Proteolytic cleavage of mucins causes disassembly of the polymerized mucin network, reducing mucus viscosity, increasing its porosity and likely impairing mucus barrier function^19, 20^. The protozoan parasite *Entamoeba histolytica* breaks down the mucus network to facilitate invasion of IECs by cleaving mucin 2 (MUC2), the major component of intestinal mucus^21^. Adult *T. muris* also degrades MUC2 via secretion of serine proteases^20^. RNA sequencing (RNA-seq) analysis of L1 larvae recovered from infected mice at 3 h and 24 h p.i. showed significantly increased expression of serine proteases (*Supplementary Fig. 3a*), as well as WAP, Kunitz and serpin classes of peptidase inhibitors (*Supplementary Fig. 3b*). It is therefore likely that L1 larvae secrete proteases to degrade mucus and this facilitates their invasion of the IE. Indeed, the sedimentation profile of purified glycosylated MUC2 was altered by exposure to L1 larvae, with a higher proportion of slower-sedimenting mucins indicating a reduction in size due to mucin depolymerization (*Fig. 3a*). Early in infections, the small ratio of larvae versus IECs in the caecum dilutes any effects the larvae have on the mucus layer; thus, from in vivo experiments, it is not possible to directly determine whether degradation occurs in the mucus layers overlaying the IECs during whipworm larvae invasion. We therefore used the caecaloid system to examine the effects of larvae on mucin more closely. Consistent with our results from purified MUC2, when comparing mucus from L1 larvae-infected and uninfected caecaloids, we observed an increased proportion of mucin distribution exhibiting a lower sedimentation rate, indicating increased degradation of mucin polymers *(Fig. 3b, Supplementary Fig. 4).* Moreover, the mucus layer immediately overlaying larvae-infected IECs was less densely stained by toluidine blue than the mucus overlaying neighbouring uninfected regions and uninfected caecaloids *(Fig. 3c)*, again indicating degradation. Collectively, these data suggest that degradation of mucus by L1 larvae enables whipworms to penetrate through the mucus layer and invade the underlying IECs.

**Figure 3.**
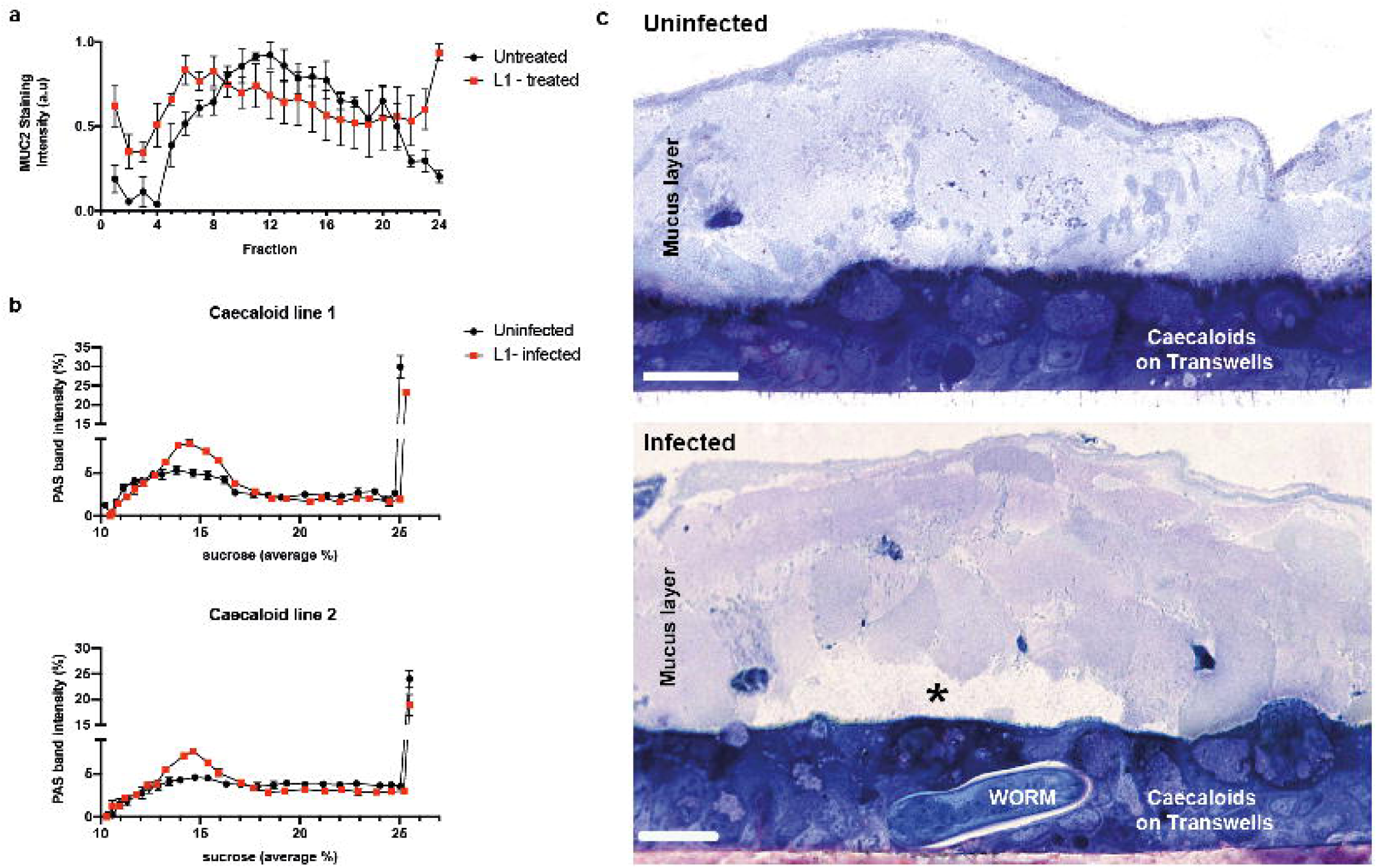
Whipworm L1 larvae invade caecal epithelium by degrading the overlaying mucus layer. **a** MUC2 purified from LS174T cell lysates was incubated with or without 400 L1 larvae at 37°C for 24 h before being subjected to rate zonal centrifugation on linear 6-8M GuHCl gradients (fraction1 low GuHCl; fraction 24 – high GuHCl). After centrifugation tubes were emptied from the top and the fractions probed with a MUC2 antibody. Data are shown as staining intensity arbitrary units (a. u). Results are represented as the mean +/-Standard Error of the Mean (SEM) of 3 independent experiments. **b** Caecaloid mucus degradation by L1 larvae at 72 h p.i. Transwells were washed with 0.2 M urea in PBS to recover mucus. Washes were subjected to rate zonal centrifugation on linear 5-25% sucrose gradients. After centrifugation tubes were emptied from the top and the fractions were stained with Periodic Acid Shiff’s (PAS) to detect the mucins. Data are shown as percentage of intensity. Results are represented as the mean +/− SEM of 3 replicas of two caecaloid lines. **c** Representative images of a toluidine blue-stained transverse sections from caecaloids uninfected and infected with *T. muris* larvae for 24 h showing degradation (asterisk) of the overlaying mucus layer immediate above the infected cells. Scale bar 20μm.

### Close interactions between *T. muris* L1 larvae and IECs defining the whipworm niche in the syncytial tunnels

After crossing the mucus layer, the L1 larva becomes intracellular creating a syncytium, a hallmark of whipworm infections *(Figs. 1* and *2)*. Only previously described for later larval and adult stages^5, 11^, syncytial tunnels are suggested to form by lateral burrowing of whipworms through adjacent IECs that join to form a single structure housing the parasite. Presently, the interactions between the host IECs and the L1 larvae and the process of formation of the tunnels are not understood. Tilney *et al.* previously observed that the syncytium around the anterior end of L3-L4 larvae and adult worms is an inert scaffold of dead cells with a brush border cover^5^. In contrast, at early stages of infection, we found that while cells left behind in the tunnel were dead, the IECs actively infected by the worm were in fact alive *(Fig. 4a).* Using TEM of caecaloids infected with *T. muris* for 24 and 72 h, we observed direct contact of the L1 larvae with the host cells cytoplasm, displacement by the larvae of cellular organelles *(Figs. 2c* and *4b-d)*, deposition of actin fibres in IECs surrounding the worm cuticle *(Fig. 4b, inset I)*, and numerous mitochondria in infected cells *(Fig 4c* and *d)*, thus recreating the host-parasite interactions observed *in vivo (Fig. 1).* With infection progression, at 72 h p.i., we also detected other alterations in infected IECs of caecaloids, including cell liquefaction and pyknotic nuclei indicating early apoptosis *(Figs. 4c* and *d)*. Moreover, several lysosomes were found in infected cells, many of which were being discharged over the larval cuticle *(Fig. 4d insets II and III).* Taken together, these findings suggest that early in infection there is an active interplay between the IECs and the parasite at its multi-intracellular niche, which may shape the initial host responses to the larvae.

**Figure 4.**
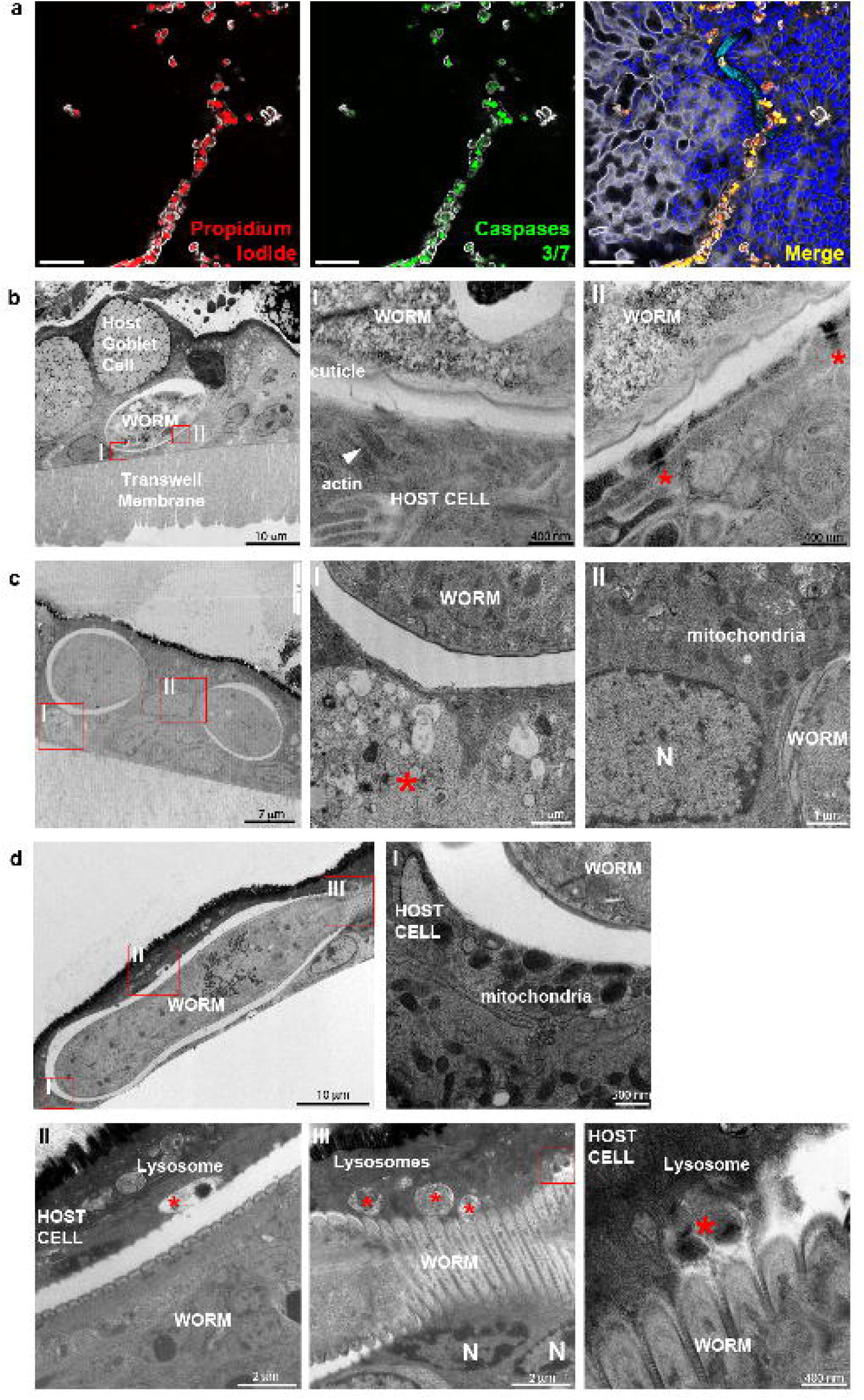
Close interactions between *T. muris* whipworm larvae and IECs at syncytial tunnels during early infection of caecaloids. **a** Selected confocal IF 2D images from a z-stack showing IECs left behind in the tunnel are necrotic (propidium iodide (red) and caspase 3/7 (green) positive), while IECs infected by worm are alive after 72h p.i. In blue and aqua, DAPI stains nuclei of IECs and larvae, respectively; and in white, Phalloidin binds to F-actin. Scale bar 50μm. **b-d** Representative TEM images of transverse sections of caecaloids infected with *T. muris* L1 larvae, showing host-parasite interactions during early infection: **b** Host cell actin fibres surround the cuticle of the worm (inset I) and desmosomes (red asterisks) are still present (inset II) at 24 h p.i. **c** Liquefied cell (inset I, asterisk), numerous mitochondria and nuclei in early stages of apoptosis (inset II) at 72 h p.i. **d** Numerous mitochondria (inset I) and lysosomes in host cells, some actively discharging over the worm cuticle (insets II and III). N, nuclei; white arrowheads, actin filaments; red asterisks, lysosomes.

### Whipworm burrowing through IECs ultimately results in tissue damage

IE barrier integrity is maintained by intercellular junctions between the IECs including, from apical to basal: tight junctions, adherens junctions, and desmosomes^22^. Syncytial tunnels hosting stage 3 and 4 larvae and adult whipworms present an intact apical surface, stabilised by the actin cytoskeleton and cell junctions, and a basal surface that remains attached to the basement membrane, but lateral membranes of the host IECs that are ruptured^5^. In contrast, during early infection, lateral membranes of host cells were still visible and separating their cytoplasm *(Figs. 1c* and *2c)* and tight junctions were still present on infected cells, but have disappeared in the cells left behind in the tunnel as indicated by the presence or absence of ZO-1 stain, in caecaloids *(Figs. 2d* and *5a, Supplementary video 5).* We noticed that while all intercellular junctions were still present in infected caecaloids cells after 72 h, desmosomes, but not tight and adherens junctions, joining infected IECs and adjacent cells had opened *(Figs. 5b, Supplementary Fig. 5a)*. When compared to those of uninfected cells, the distance between desmosomes joining infected IECs and adjacent cells significantly increased from 26 nm to 38 nm *(Fig. 5b)*. Strikingly, we observed equivalent perturbations *in vivo (Supplementary Fig. 5)*, further demonstrating that the caecaloid-whipworm model closely recapitulates whipworm infection. Altogether, our results indicate that with progression of infection, the tunnelling of the larvae through the IECs results in IEC damage.

**Figure 5.**
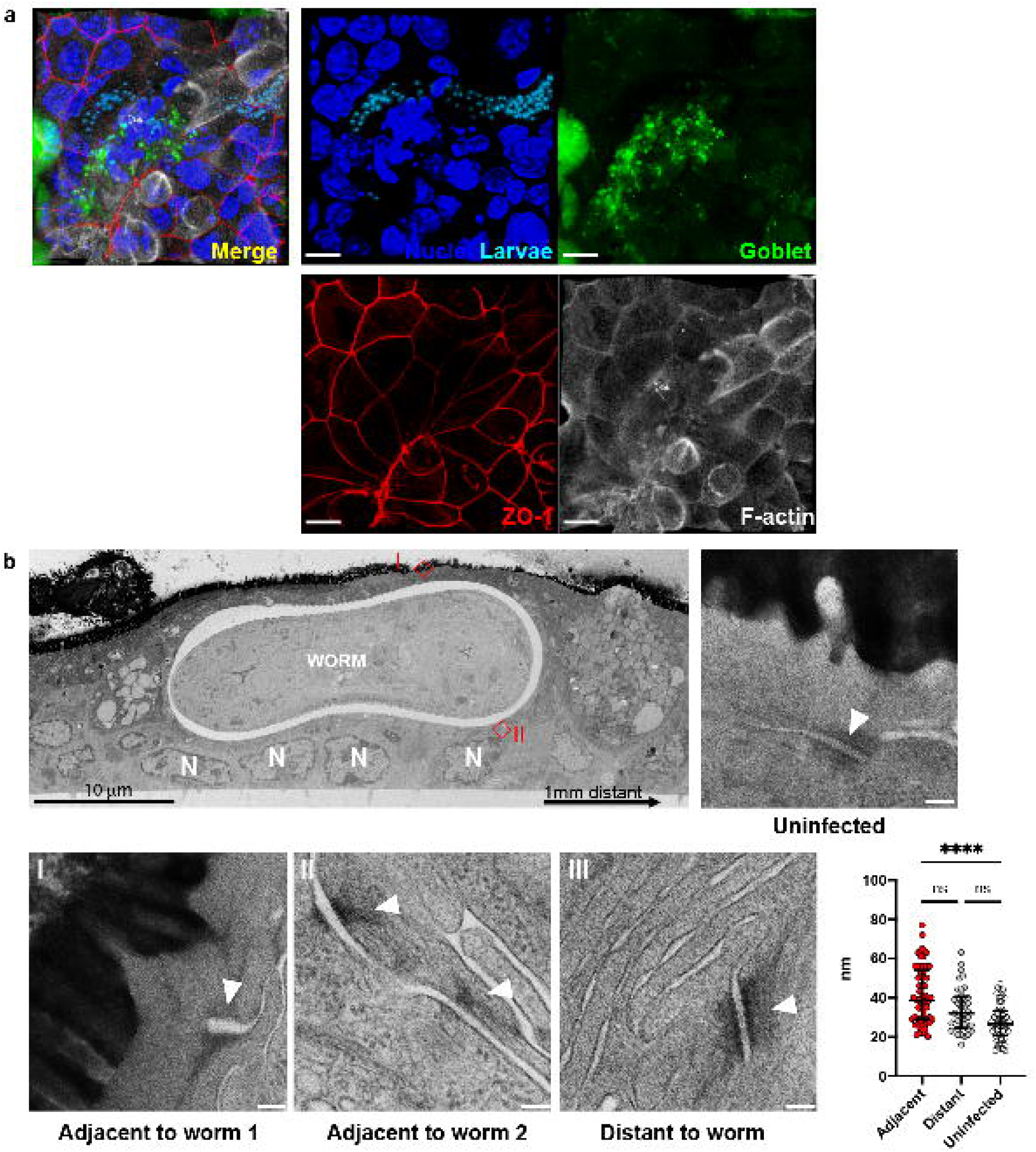
Perturbations on desmosomes, but not in tight junctions, in host cells of whipworm larvae during early infection of caecaloids. **a** Z-stack projection of confocal IF images of larva in syncytial tunnel in caecaloids infected with L1 whipworm larvae for 24 h. In red, ZO-1 protein binds tight junctions; in green, the lectins UEA and SNA bind mucins in goblet cells; in blue and aqua, DAPI stains nuclei of IECs and larvae, respectively; and in white, Phalloidin binds to F-actin. Scale bar 10μm. **b** Representative TEM images of transverse sections of *T. muris-*infected caecaloids (72 h p.i.) and desmosomes (arrowheads) joining infected and adjacent cells (insets I and II), cells 1mm distant to the worm from infected caecaloids (inset III), and cells from uninfected caecaloids. Scale bars for desmosome images 100nm. Desmosome separation in nm was measured in host cells from four independent worms. Measurements adjacent n=62, distant n=37 and uninfected n=50. ****p<0.0001 Kruskal Wallis test and Dunn’s comparisons among groups.

### Host IECs responses to early infection with whipworms are dominated by a type-I IFN signature

IEC responses to whipworm early infection are thought to initiate host immune responses to the worm and orchestrate repair to the IE damage caused by larval invasion and tunnelling^13, 14^. However, currently little is known about the nature of those responses. Using bulk RNA-seq of whole caeca from infected mice at day 7 p.i and caecal IECs from infected mice at 24 and 72 h p.i., we detected the upregulation of genes involved in innate immune responses, specifically those related to type-I IFN signalling and normally characteristic of bacterial and viral infections *(Fig. 6a, Supplementary Fig. 6, Supplementary Data 1* and *2)*. The response to larval infection in the caecum at day 7 p.i. therefore appeared to be driven by the IECs from a much earlier stage of infection.

**Figure 6.**
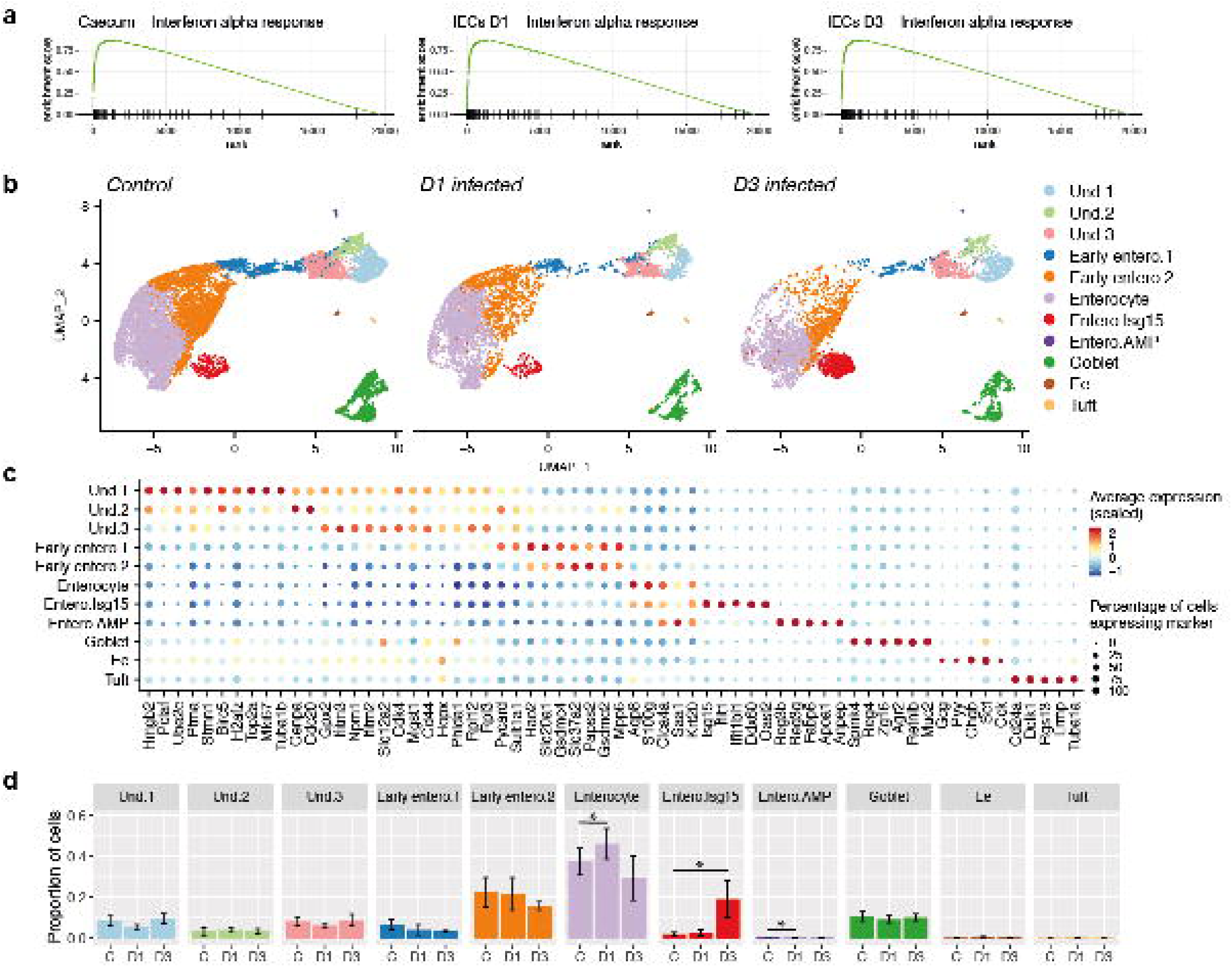
Host IECs responses to early infection with whipworms are dominated by a type-I IFN signature. **a** Bulk RNA-seq data from complete caecum and caecal IECs of *T. muris*-infected and uninfected mice at days 7, and 1 and 3 p.i., respectively, were analysed by gene set enrichment analysis (GSEA) for cell signature genes in the IFN alpha pathway. All analyses have false discovery rate (FDR) adjusted p values: Caecum, 0.013; day 1 (D1) IEC, 0.025; day 3 (D3) IEC, 0.026. **b** Uniform manifold approximation and projection (UMAP) plots from single cell RNA-seq analysis of 22422 EpCAM^+^CD45^-^ cells. IEC populations (colour coded) in the caecum of control (n=8, 4 mice for each time point) and *T. muris*-infected mice after 1 and 3 days p.i. (n=4 mice for each time point). **c** Dot plot of the top marker genes for each cell type. The relative size of each dot represents the fraction of cells per cluster that expresses each marker; the colour represents the average (scaled) gene expression. **d** Increased relative abundance of the Enterocyte Isg15 cluster upon 72 h of *T. muris* infection. The size of the clusters, expressed as a proportion of the total number of cells per individual, was compared across four biological replicates at each time point for uninfected and *T. muris-*infected mice. Mean +/− standard deviation is shown (* p<0.05, two-tailed t test).

Using the 10X Chromium platform, we performed single-cell RNA-seq (scRNA-seq) of caecal IECs from uninfected and *T. muris*-infected mice at 24 and 72h p.i. Populations of undifferentiated, enterocytes, goblet, enteroendocrine and tuft cells could be identified (*Fig. 6b-d, Supplementary Fig. 7*). Isolating undifferentiated cells *in silico*, we further characterised 5 subpopulations: stem and TA cells that are on the S and G2/M phases of cell cycle, deep secretory cells and two enterocyte progenitor populations, which express known markers of these cell types in the small intestine^23, 24^ and colon^25–29^ (*Supplementary Figs. 8* and *9*). Enterocytes were divided in five sub-clusters: two early enterocyte populations and three late/mature ones, distinguished by the expression of particular marker genes involved in defined biological processes (*Figs. 6b* and *c, Supplementary Data 3*.). Interestingly, one cluster of enterocytes was characterised by the expression of IFN-stimulated genes (ISGs), specifically *Isg15, Ifit1, Ifitbl1, Ddx60* and *Oasl2* (*Fig. 6b* and *c*). We detected a striking increase in the size of this cluster on *T. muris*-infected mice at 72 h p.i. *(Figs. 6b* and *d)*.

We validated the expansion of *Isg15* expressing enterocytes in response to whipworm infection using mRNA *in situ* hybridization (ISH) by chain reaction (HCR) on caecal tissues from uninfected and *T. muris*-infected mice after 24 and 72 h p.i. *(Fig. 7a-c)*. In uninfected mice, occasional crypts expressing high levels of *Isg15* were detected *(Fig. 7a)*, consistent with the presence of this cluster of enterocytes in naïve mice *(Fig. 6b).* These crypts were easily distinguishable above the extremely low baseline level of *Isg15* expressed in enterocytes throughout the caecum. Interestingly, by 72h p.i., the number of crypts showing high levels of *Isg15* expression increased significantly, with those *Isg15^+^* crypts forming large groups or “islands” *(Figs. 7b-c)*. *T. muris* is detectable *in situ* by its expression of p43 *(Figs. 7d-e)*, the single most abundant protein secreted/excreted by the parasite^30^. Using multiplexed ISH by HCR for *T. muris p43*, mouse *Krt20* (mature enterocyte marker, *Fig 6c*) and *Isg15*, we were able to locate worms in the caecum of infected mice. Larvae were occasionally found near islands of *Isg15^+^* crypts (*Fig. 7f*; n=3/9 worms detected at 72h p.i.), though this was not always the case (*Fig. 7g*; n=6/9 worms detected at 72h p.i.). We speculate that larval infection and tunnelling through IECs at the bottom of the crypts resulted in the activation of responses by enterocytes immediately above. Taken together, these findings suggest that host IECs responses to early whipworm larvae infection are dominated by a type-I IFN signature driven by the expansion of a distinct population of enterocytes expressing *Isg15*.

**Figure 7.**
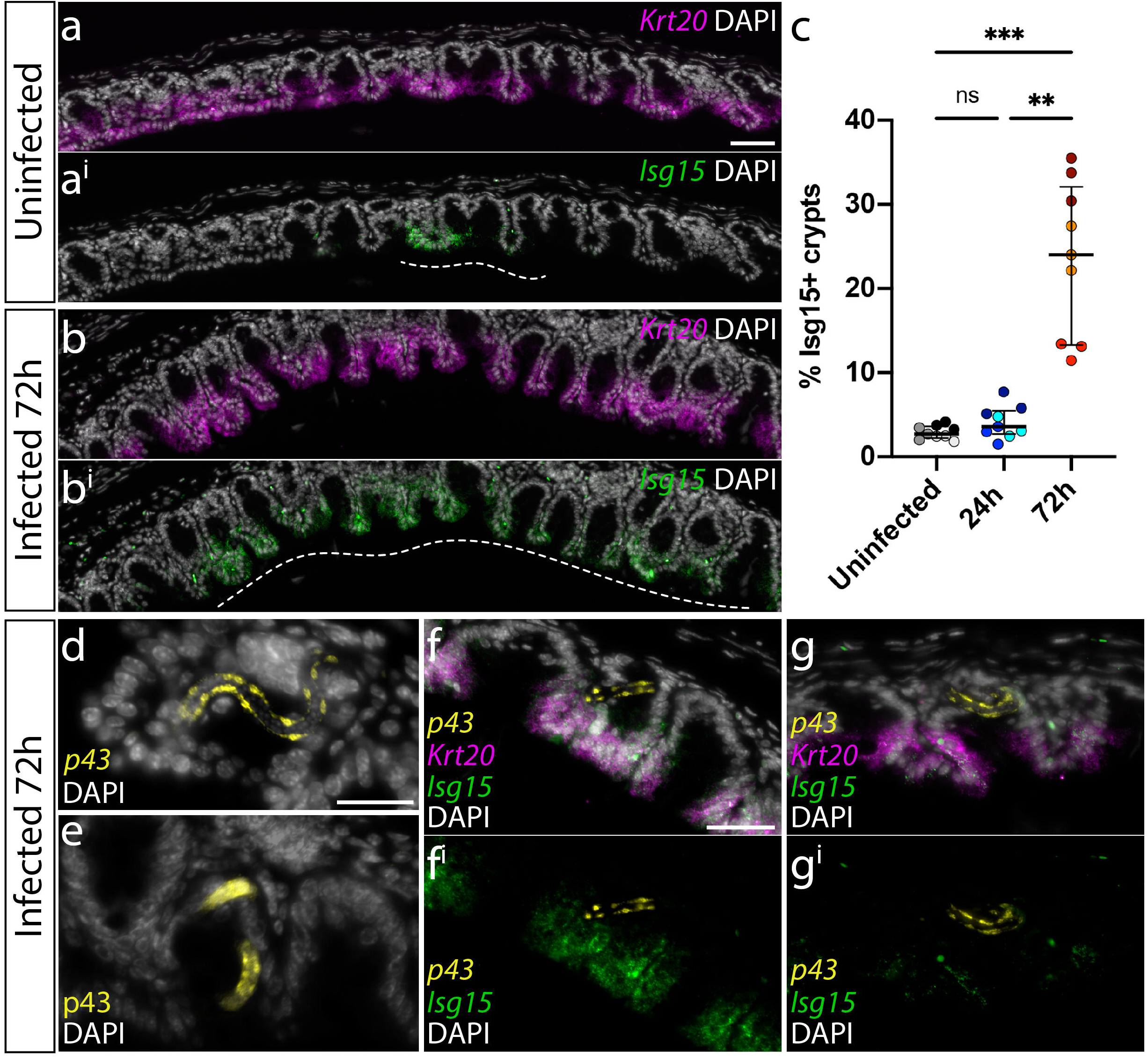
Expansion of crypts with enterocytes expressing *Isg15* upon whipworm infection. Representative images of expression of *Isg15* (green) and *Krt20* (magenta), visualised by mRNA ISH by HCR, in the caecum of **a** uninfected mice and **b** *T.muris*-infected mice after 72 h of infection. Dashed white lines show the extent of “islands” of *Isg15^+^* crypts. **c** The number of *Isg15^+^* crypts in a caecal section was calculated as a percentage of the total number of crypts. Three caecal sections (technical replicates) per mouse were quantified, with three mice analysed per condition (uninfected, 24 and 72 h p.i.), **p=0.0045, ***p=0.0002 Kruskal Wallis test and Dunn’s comparisons among groups. For each condition, dots representing technical replicates are coloured identically. **d** Detection of *T. muris p43* transcripts (by HCR) and **e** p43 protein (by IF) facilitated location of worms within infected caecal tissue. **f** In some instances, worms were located near islands of *Isg15^+^* enterocytes, **g** while in other cases, worms were found away from these islands. Scale bars: a/a^i^ and b/b^i^=60 μm; d and e = 25 μm; f/f^i^ and g/g^i^=30 μm.

## Discussion

We have shown that whipworm invasion of the IE is preceded by the degradation of the mucus layers and identified absorptive and goblet cells as the main constituents of the syncytium hosting whipworm larvae. Our findings revealed the early syncytial tunnels are an interactive multi-intracellular niche where whipworms interplay with their host IECs resulting in the activation of type-I IFN signalling responses that potentially orchestrate the development of immunity against the worm and tissue repair.

Previous studies on syncytial tunnels have focused on L3-L4 larval and adult stages when, in the caecal and proximal colonic crypts, the syncytium becomes a visible protrusion of the wall into the lumen^5, 11^. In this work, we determined interactions between *T. muris* L1 larvae and IECs mediating whipworm penetration of the caecal epithelium and formation of syncytial tunnels. To do this, we exploited a novel *in vitro* system that used caecaloids, together with *in vivo* observations where possible. Caecaloids grown and differentiated in transwells recapitulate the complexity of the cellular composition and, to some degree, the architecture of the caecal epithelium. This architecture promotes whipworm larvae infection *in vitro* enabling us to tackle questions that could not otherwise be investigated in the mouse model. The fact that whipworm invasion and colonisation is not supported by epithelial cell lines, 3D caecaloids or undifferentiated caecaloids grown in transwells (forming a complete monolayer but lacking differentiated cell types) indicates that a combination of interactions of the larvae with particular cellular or molecular components of the caecal epithelium, as well as specific physicochemical conditions, are critical in triggering parasite invasion. Those components include a complex mucus layer that is well mimicked by our *in vitro* system, which allowed us to visualise mucus degradation *in situ* and detect depolymerisation of mucins upon larvae infection. Other pathogens that preferentially colonise the large intestine such as *Shigella dysenteriae*^31^ and *E. histolytica*^32, 33^ recognize tissue specific expression of mucin and mucin glycosylation patterns. *Trichuris* L1 larvae may also recognise molecular cues within the mucus to initiate invasion as Hasnain *et al.* showed decreased establishment of *T. muris* in *Sat1^-/-^* mice with reduced sulphated mucus^34^. For penetration and tunnelling through IECs, the expression of proteases by whipworm L1 larvae we detected may be critical as proteases secreted by other parasitic nematode infective larvae facilitate their entry in the intestinal wall of mammalian hosts^35, 36^. Future studies we be able to use the caecaloid-whipworm larvae model together with live imaging tools to capture active invasion, and knock-out technologies and inhibitors, will determine the nature of the molecular mechanisms directing larvae movement towards and penetration at the IECs of the crypt base.

Our findings on the cellular composition of early tunnels are in line with previous observations indicating that absorptive and goblet cells constitute the syncytium hosting older larval stages and adults^11^. Ascertaining the path of the L1 larvae inside the IECs of caecaloids allowed us to discover that while the IECs ultimately succumb after the parasite has moved through them, when the larvae are within the IECs, they remain alive. Surprisingly, during the first 24 h of infection these host cells present minimal damage. Despite the presence of larva in direct contact with their cytoplasm, which has joined into a multinucleated mass, desmosomal contacts with adjacent cells and lateral membranes are conserved. We identified distinct host-parasite interactions in the tunnels including the reorganisation of the infected IEC cytoskeleton around the cuticle of the larvae suggesting a response of the host cell to the intracellular parasite. Moreover, we visualised a likely excretion/secretion of products by the larva into its immediate environment in the cytoplasm of the infected IECs. These products may support larval intracellular tunnelling by digesting cells and cellular components, or manipulating the activation of inflammatory responses by the IECs. We also detected numerous mitochondria in infected IECs when compared with uninfected neighbouring cells. Within early syncitia, larvae may therefore interact with their host cells to modify their energy demands and metabolic activity. As infection progressed, at 72 h p.i., we identified infected cells where several lysosomes had surrounded and discharged over the cuticle of the larvae, suggesting a direct response of the host cell to the parasite. In addition, we observed perturbations in IECs forming the syncytial tunnel including opening of the desmosomes, early pyknotic nuclei indicative of apoptosis and liquefaction of cells. These alterations indicate that whipworm burrowing through IECs ultimately results in IE damage.

IE damage at the syncytial tunnels caused by the whipworm larvae either mechanically through tunnelling or chemically as induced by secretory/excretory products, potentially results in release of damage-associated molecular patterns (DAMPs) and alarmins by infected IECs, which have been suggested to initiate innate immune responses to the worm^13, 14, 37^. The production of the alarmins IL-25^38, 39^, IL-33^40, 41^ and TSLP^42, 43^ implicated in the induction of type 2 responses to *Trichuris* and other parasitic worms^37^ was either not detected or did not significantly increase in our single cell or bulk transcriptomics data over the first 72 hours of infection *(Supplementary Fig. 10 and 11).* It is remarkable that upon invasion of the epithelium over the very early stages of whipworm colonisation, IEC responses to the worm are mostly silent. Conversely, in response to infection we identified a striking expansion of an enterocyte population expressing several ISGs including *Isg15*. ISG15 has been described as an alarmin that induces tissue alerts and inflammation via immune cell recruitment, infiltration and activation^15–17^. On the other hand, ISG15 exerts immunoregulatory functions by negatively regulating type-I IFN signalling and production of proinflammatory cytokines and chemokines^16, 44^. Moreover, ISG15 regulates tissue damage and/or repair responses, specifically in the context of viral infection of the respiratory epithelium^16, 45^. Intestinal, respiratory and corneal epithelial cells have been shown to express *Isg15* in response to not only viral, but also protozoal parasitic and fungal infections^16, 46–48^ and to inflammation during IBD^17^. We therefore hypothesise that ISG15 acts as an alarmin released upon of IECs damage caused by whipworm larvae invasion, potentially triggering the development of immune and tissue repair responses.

ISG15 is rapidly induced in response to type-I IFNs but also to nucleic acids sensed by cytosolic receptors in an IFN-independent manner^16^. Our transcriptomic data suggests that the larval induction of *Isg15* in IECs is type-I IFN independent. While we did not detect increased levels of *Ifna1* and *Ifnab1* upon infection either in the complete caecum or IECs *(Supplementary Fig. 10 and 11)*, the expanded *Isg15*-expressing enterocyte population co-expressed *Ddx60* and *Irf7*. DDX60 promotes retinoic-acid inducible gene I (RIG-I)-like receptor mediated signalling upon cytosolic sensing of self- and non-self nucleic acids that results in *Isg15* expression via the IFN regulatory factors 3 and 7 (IRF3, IRF7)^16, 49^. Thus, DNA or RNA released from whipworm infected/damaged IECs or secreted by larvae may induce *Isg15* expression in infected and bystander cells. Further research is needed to understand the molecular mechanisms leading to *Isg15* expression by IECs and role of this alarmin in host responses to whipworm infection.

Studies on infection with other parasitic worms, including *Schistosoma mansoni*^50^, *Nippostrongylus brasiliensis*^51^ and *Heligmosomoides polygyrus*^52^ suggest that type-I IFNs are important in driving the initiation of type 2 responses that result in worm expulsion^37^. Our data indicates that this is not the case for *T. muris*, which would be of clear benefit to the parasite allowing it to remain in its niche, and furthermore exemplifies the diverse regulatory mechanisms that govern early host responses to different helminths.

Collectively, our work has contributed to define the early stages of intestinal invasion and colonisation by whipworms. Extending the applicability of our caecaloid-whipworm system, adapting it to study *T. trichiura* L1 larvae infection of human caecal and proximal colon organoids and adding stroma and immune cells will be important next steps. Further investigations on the early host-parasite interplay within the whipworm mucosal niche will be fundamental to the development of new tools to help control trichuriasis and also provide novel insights into how the intestinal epithelium adapts to damage and mediates repair.

## Methods

### Mice

C57BL/6N mice were kept under specific pathogen-free conditions, and colony sentinels tested negative for *Helicobacter* spp. Mice were fed a regular autoclaved chow diet (LabDiet) and had *ad libitum* access to food and water. All efforts were made to minimize suffering by considerate housing and husbandry. Animal welfare was assessed routinely for all mice involved. Mice were naïve prior the studies here described. Experiments were performed under the regulation of the UK Animals Scientific Procedures Act 1986 under the Project licenses 80/2596 and P77E8A062 and were approved by the institutional Animal Welfare and Ethical Review Body.

### Parasites and *T. muris* infection

Infection and maintenance of *T. muris* was conducted as described^53^. Age and sex matched female mice (6–10-week-old) were orally infected under anaesthesia with isoflurane with a high (400-1000) dose of embryonated eggs from *T. muris* E-isolate. Mice were randomised into uninfected and infected groups using the Graph Pad Prism randomization tool. Uninfected and infected mice were co-housed. Mice were monitored daily for general condition and weight loss. Mice were culled including concomitant uninfected controls at different time points by cervical dislocation, and caecum and proximal colon were collected for downstream processing. Blinding at the point of measurement was achieved using barcodes. During sample collection, group membership could be seen, however this stage was completed by technician staff with no knowledge of the experiment objectives.

### In vitro hatching of T. muris eggs with Escherichia coli

*E. coli* K-12 was grown in Luria Bertani broth overnight at 37°C and shaking at 200 rpm. Eggs were added to bacterial cultures and incubated for 2 h at 37°C, 5% CO_2_. Larvae were washed with PBS three times to remove *E. coli* by centrifugation at 720 *g* for 10 min at RT. Bacteria were killed by culturing larvae in RPMI 1640 (Gibco Thermo Fisher Scientific), 10% Foetal Bovine Serum (FBS) (Gibco Thermo Fisher Scientific), 2 mM L-glutamine (Sigma-Aldrich), 1X antibiotic/antimycotic (Sigma Aldrich) and 1 mg/ml ampicillin (Roche) for 2 h at 37°C, 5% CO_2_. Larvae were washed with RPMI 1640 three times to remove ampicillin and separated from egg shells and unembryonated eggs using a stepped 50-60% Percoll (Sigma Aldrich) gradient. Centrifugation at 300 *g* for 15 min at RT was performed and the 50% interface layer was collected. Recovered larvae were washed with RPMI 1640 and resuspended in media containing Primocin (InvivoGen).

### *T. muris* L1 and L2 larvae recovery from infected mice for RNA extraction

Mice were culled at 3 and 24 h post infection (p.i). to recover L1 larvae and at day 14 to recover L2 larvae. Caecum and proximal colon were collected and placed in 5X penicillin/streptomycin (Gibco Thermo Fisher Scientific) in Dulbecco’s PBS 1X without calcium and magnesium (PBS) (Gibco Thermo Fisher Scientific). Caecum and proximal were cut longitudinally and were washed to remove faecal contents. The tissues were cut into small sections and added to 0.9% NaCl in PBS and incubated in a water bath at 37°C for 2 h to allow L1/L2 larvae to come free from the epithelium. Larvae were removed from the NaCl and placed into 1X penicillin/streptomycin PBS. Larvae were washed once with PBS and pellets were resuspended in TRIzol LS (Invitrogen).

### 3D Caecaloid culture

Mouse 3D caecaloids lines from C57BL/6N adult mice (6-8 weeks old) were derived from caecal epithelial crypts as previously described^18^. Briefly, the caecum was cut open longitudinally and luminal contents removed. Tissue was then minced, segments were washed with ice-cold PBS and vigorous shaking to remove mucus, and treated with Gentle Cell Dissociation Reagent (STEMCELL Tech) for 15 min at room temperature (RT) with continuous rocking. Released crypts were collected by centrifugation, washed with ice-cold PBS, resuspended in 200 μl of cold Matrigel (Corning), plated in 6-well tissue culture plates and overlaid with a Wnt-rich medium containing base growth medium (Advanced DMEM/F12 with 2 mM Glutamine, 10 mM HEPES, 1X penicillin/streptomycin, 1X B27 supplement, 1X N2 supplement (all from Gibco Thermo Fisher Scientific)), 50% Wnt3a-conditioned medium (Wnt3a cell line, kindly provided by the Clevers laboratory, Utrecht University, Netherlands), 10% R-spondin1 conditioned medium (293T-HA-Rspo1-Fc cell line, Trevigen), 1 mM N-acetylcysteine (Sigma-Aldrich), 50 ng/ml rmEGF (Gibco Thermo Fisher Scientific), 100 ng/ml rmNoggin (Peprotech), 100 ng/ml rhFGF-10 (Peprotech) and 10 μM Rho kinase (ROCK) inhibitor (Y-27632) dihydrochloride monohydrate (Sigma-Aldrich). Caecaloids were cultured at 37°C, 5% CO_2_. The medium was changed every two days and after one week, Wnt3a-conditioned medium was reduced to 30% and penicillin/streptomycin was removed (expansion medium). Expanding caecaloids were passaged, after recovering from Matrigel using ice-cold PBS or Cell Recovery Solution (Corning), by physical dissociation through vigorous pipetting with a p200 pipette every six to seven days.

### Caecaloid culture in 2D conformation using transwells

3D caecaloids grown in expansion medium for 4-5 days after passaging were dissociated into single cells by TrypLE Express (Gibco Thermo Fisher Scientific) digestion. 200,000 cells in 200 µl base growth medium were seeded onto 12 mm transwells with polycarbonate porous membranes of 0.4 µm (Corning) pre-coated with 50 mg/ml rat tail collagen I (Gibco Thermo Fisher Scientific). Cells were cultured with expansion medium in the basolateral compartment for two days. Then, basolateral medium was replaced with medium containing 10% Wnt3a-conditioned medium for additional 48 h. To induce differentiation of cultures, medium in the apical compartment was replaced with 50 µl base growth medium and medium in the basolateral compartment with medium containing 2.5% Wnt3A-conditioned medium that was changed every two days. Cultures were completely differentiated when cells pumped the media from the apical compartment and cultures looked dry.

### *T. muris* L1 larvae infection of caecaloids grown in transwells

Differentiated caecaloid cultures in transwells were infected with 300 L1 *T. muris* larvae obtained by *in vitro* hatching of eggs in presence of *E. coli*. Larvae in a volume of 100 μl of base growth medium were added to the apical compartment of the transwells. Infections were maintained for up to 72 h at 37°C, 5% CO_2_.

### IF staining of caecaloids

For IF, caecaloid cultures in transwells were fixed with 4% Formaldehyde, Methanol-free (Thermo Fisher) in PBS for 20 min at 4°C, washed three times with PBS and permeabilized with 2% Triton X-100 (Sigma-Aldrich) 5% FBS in PBS for 1 h at RT. Caecaloids were then incubated with primary antibodies α-villin (1:100, Abcam, ab130751), α-Ki-67 (1:250, Abcam, ab16667), α-chromogranin A (1:50, Abcam, ab15160), α-Dcamlk-1 (1:200, Abcam, ab31704), α-zona occludens-1 (ZO-1) protein (1:200, Invitrogen, 61-7300) and the lectins *Ulex europaeus* agglutinin - Atto488 conjugated (UEA, 1:100, Sigma-Aldrich, 19337) and *Sambucus nigra -* Fluorescein conjugated (SNA, 1:50, Vector Laboratories, FL-1301) diluted in 0.25% Triton X-100 5% FBS in PBS overnight at 4°C. After three washes with PBS, caecaloids were stained with secondary antibody Donkey anti-rabbit IgG Alexa Fluor 555 (1:400, Molecular Probes, A31572), phalloidin Alexa Fluor 647 (1:1000, Life Technologies, A22287) and 4’,6’-diamidino-2-phenylindole (DAPI, 1:1000, AppliChem, A1001.0010) at RT for 1 h. Transwell membranes were washed three times with PBS and mounted on slides using ProLong Gold anti-fade reagent (Life Technologies Thermo Fisher Scientific). Confocal microscopy images were taken with a Leica SP8 confocal microscope and processed using the Leica Application Suite X software.

### Cell death fluorescence staining of caecaloids

To evaluate cell death in infected caecaloids, cultures were incubated with 100 μl warm base growth medium containing 0.3 mg/ml of propidium iodide (Sigma-Aldrich) and 8 μM of CellEvent™ Caspase-3/7 Green Detection Reagent (Invitrogen) for 30 min at 37°C 5% CO2. Then, caecaloids were fixed and counterstained as described above.

### *mRNA* ISH by HCR and IF on paraffin sections of mice

Caeca from uninfected and *T. muris*-infected mice after 24 and 72 h p.i. were fixed, embedded in paraffin and sectioned at 8 µm thickness for mRNA *in situ* hybridization as previously described^54^. All probes, buffers, and hairpins for third generation HCR were purchased from Molecular Instruments (Los Angeles, California, USA). HCR on paraffin sections was carried out according to the protocol of Choi et al. (2018)^55^, with modifications according to Criswell and Gillis (2020)^56^.

Immunofluorescence on paraffin sections was carried out according to the protocol of Marconi *et al.* (2020)^54^, except with heat-mediated antigen retrieval. Antigen retrieval was performed by warming dewaxed and rehydrated slides in water at 60°C for 5 min, followed by incubation in citrate buffer (10 mM sodium citrate, 0.05% Tween20, pH 6.0) at 95°C for 25 min. Slides were then cooled for 30 min at −20°C and rinsed 2 x 5 min in 1X PBS + 0.1% Triton X-100 at RT before proceeding with blocking and antibody incubation. Rabbit anti-p43 primary and AF488-conjugated goat-anti-rabbit IgG secondary antibodies were diluted 1:400 and 1:500, respectively, in 10% sheep serum. All slides were coverslipped with Fluoromount-G containing DAPI (Southern Biotech) and imaged on a Zeiss Axioscope A1 compound microscope.

### Transmission EM

Caeca and caecaloids were fixed in 2.5% glutaraldehyde/2% paraformaldehyde in 0.1M sodium cacodylate buffer, post-fixed with 1% osmium tetroxide and mordanted with 1% tannic acid followed by dehydration through an ethanol series (contrasting with uranyl acetate at the 30% stage) and embedding with an Epoxy Resin Kit (all from Sigma-Aldrich). Semi-thin 0.5 μm sections were cut and collected onto clean glass slides and dried at 60°C before staining with 1% Toluidine Blue and 1% Borax (all from Sigma-Aldrich) in distilled water for 30 seconds. Sections were then rinsed in distilled water and mounted in DPX (Sigma-Aldrich) and coverslipped. Sections were imaged on a Zeiss 200M Axiovert microscope.

Ultrathin sections cut on a Leica UC6 ultramicrotome were contrasted with uranyl acetate and lead nitrate, and images recorded on a FEI 120 kV Spirit Biotwin microscope on an F416 Tietz CCD camera.

### Scanning EM

Caecaloids were fixed with 2.5% glutaraldehyde and 4% paraformaldehyde in 0.01 M PBS at 4°C for 1 h, rinsed thoroughly in 0.1 M sodium cacodylate buffer three times, and fixed again in 1% buffered osmium tetroxide for 3 h at RT. To improve conductivity, using the protocol devised by Malick and Wilson^57^, the samples were then impregnated with 1% aqueous thiocarbohydrazide and osmium tetroxide layers, with the steps separated by sodium cacodylate washes. They were then dehydrated three times using an ethanol series (30, 50, 70, 90, and 100% ethanol, 20 min each) before they were critical point dried in a Leica EM CPD300 and mounted on aluminium stubs with conducting araldite and sputter coated with a 2 nm platinum layer in a Leica EM ACE 600. Images were taken on a HITACHI SU8030.

### Serial block face scanning EM

Samples from caeca of infected mice were processed according to Deerinck et al^58^. Embedded tissues were mounted and serial sectioned on a Gatan 3View System and simultaneously imaged on a Zeiss Merlin SEM. Serial images were oriented and assimilated into corrected z-stacks using IMOD. The phenotype of infected cells was determined in each image and number cells of each cellular population were quantified.

### MUC2 and caecaloid mucus degradation experiments

Glycosylated MUC2 was derived from LS174T cells, purified as previously described^20^ and incubated with 400 *T. muris* L1 larvae for 24 h at 37°C, 5% CO_2_. For experiments with caecaloid cultures, after 24 and 72 h of L1 larvae infection, mucus was recovered by six PBS washes followed by six washes with 0.2 M urea.

### Rate zonal centrifugation

Mucus degradation analysis was conducted as described^20^. Briefly, purified MUC2 (in 4 M guanidinium hydrochloride (GuHCl)) was loaded onto the top of 6–8 M GuHCl gradients and centrifuged in a Beckman Optima L-90K Ultracentrifuge (Beckman SW40 rotor) at 40,000 rpm for 2.75 h 15°C. Alternatively, mucus samples (in PBS or 0.2 M urea) were loaded onto 5-25% (w/v) linear sucrose gradients and centrifuged in a Beckman Optima L-90K Ultracentrifuge (Beckman SW40 rotor) at 40,000 rpm for 3 h 15°C. After, centrifugation tubes were emptied from the top and the fractions probed with a MUC2 antibody or using the Periodic Acid Shiff’s (PAS) assay. To determine the sucrose or GuHCl concentration the refractive index of each fraction was measured using a refractometer; all sucrose and GuHCl gradients were comparable (data not shown).

### Caecal IECs Isolation

Caeca of uninfected and infected mice at day one and three p.i. were processed individually in parallel. Caeca were open longitudinally, washed with ice-cold HBSS 1X (Gibco Thermo Fisher Scientific) containing 1X penicillin/streptomycin to remove the caecal contents and cut in small fragments. These were incubated at 37°C in DMEM High Glucose (Gibco Thermo Fisher Scientific), 20% FBS, 2% Luria Broth, 1X penicillin/streptomycin, 100 µg/ml gentamicin (Sigma Aldrich), 10 µM ROCK inhibitor and 0.5 mg/ml Dispase II (Sigma Aldrich) with horizontal shaking for 90 min to detach epithelial crypts. Supernatant containing crypts was filtered through a 300 µm cell strainer (PluriSelect) and pelleted by centrifugation at 150 *g* for 5 min at RT. Crypts were dissociated into single cells by TrypLE Express digestion 10-20 min at 37°C. The epithelial single-cell suspension was washed and counted using MOXI automated cell counter. Cells were stained using the antibodies anti-CD236 (epithelial cell adhesion molecule (EPCAM); PE-Cy7, Biolegend) and anti-CD45 (Alexa 700, Biolegend) for 20 min on ice. Cells were washed and stained with DAPI. Live epithelial cells (CD236+, CD45-, DAPI-) were sorted using a fluorescence-activated cell sorting (FACS) Aria flow cytometer (BD) into TRIzol LS for bulk-RNAseq or DMEM High Glucose 10% FBS 10 µM ROCK inhibitor for droplet-based single-cell RNAseq.

### Bulk RNA isolation from L1 and L2 *T. muris* larvae and library preparation for RNA-seq

L1/L2 larvae were collected via pipette in 200-300 µl of PBS then added directly to lysing matrix D (1.4-mm ceramic spheres, MagNA Lyser Green Bead tubes, Roche) along with 1 ml of TRIzol LS. A PreCellys 24 (Bertin) was used to homogenize the samples by bead beating 3x for 20 sec at 6000 rpm, placing the samples on wet ice to cool between runs. Homogenized samples were transferred to 1.5 ml microfuge tubes and total RNA extracted by adding 400 µl of Chloroform, shaking vigorously and incubating for 5 min at RT. Samples were centrifuged at 12000 *g* for 15 min at RT, then the upper aqueous phase was transferred to a new microfuge tube prior to addition of 800 µl Isopropyl alcohol with 2 µl GlycoBlue (Invitrogen). Tubes were placed at −80 °C overnight and then centrifuged at 12000 *g* for 10 min at 4 °C. The supernatant was removed, being careful not to disturb the blue pellet, and washed in 1 ml 75 % ethanol. The pellet was air-dried briefly then resuspended in nuclease-free water. Total RNA was quantified by Bioanalyzer (Agilent). Multiplexed cDNA libraries were generated from 300 pg of total high-quality RNA according to the SmartSeq2 protocol by Picelli et al. (2014)^59^, and 125bp paired-end reads were generated on the Illumina HiSeq according to the manufacturer’s standard sequencing protocol.

### Bulk RNA isolation from caecum of uninfected and *T. muris*-infected mice and library preparation for RNA-seq

Total RNA from sections of caecum of mice pre-infection, and 4 and 7 days p.i., was isolated using 1 ml TRIzol and lysing matrix D to homogenise tissues with a Fastprep24 (MP Biomedicals), and following manufacturer’s standard extraction protocol. Total RNA was quantified by Bioanalyzer and 1 µg or 50µl cherry picked. Poly A mRNA was purified from total RNA using oligodT magnetic beads and strand-specific indexed libraries were prepared using the Illumina’s TruSeq Stranded mRNA Sample Prep Kit followed by ten cycles of amplification using KAPA HiFi DNA polymerase (KAPA Biosystems). Libraries were quantified and pooled based on a post-PCR Bioanalyzer and 75 base pair (bp) paired-end reads were generated on the Illumina HiSeq 2500 according to the manufacturer’s standard sequencing protocol.

### Bulk RNA isolation from sorted IECs from uninfected and *T. muris*-infected mice and library preparation for RNA-seq

Total RNA from sorted IECs 1 and 3 days p.i. with time matched uninfected concomitant controls, was isolated using Trizol LS. Briefly, 200 µl of Chloroform was added to 500 µl of samples in Trizol LS, shaked vigorously and incubated for 5 min at RT. Samples were centrifuged at 15000 *g* for 15 min at RT and the upper aqueous phase was recovered and mixed with one volume of 100% ethanol. RNA was recovered using the RNA Clean and Concentrator kit (Zymo Research). The samples were quantified with the QuantiFluor RNA system (Promega) and 100 ng/50 µl cherry picked. Libraries were then constructed using the NEB Ultra II RNA custom kit (New England BioLabs) on an Agilent Bravo WS automation system followed by 14 cycles of PCR using KAPA HiFI Hot Start polymerase (KAPA Biosystems). The libraries were then pooled in equimolar amounts and 75bp paired-end reads were generated on the Illumina HiSeq 4000 according to the manufacturers standard sequencing protocol.

### Droplet-based single-cell RNA sequencing 10X

Sorted cells were counted using MOXI automated cell counter and loaded onto the 10X Chromium Single Cell Platform (10X Genomics) at a concentration of 1000 cells per µl (Chromium Single Cell 3’ Reagent kit v.3) as described in the manufacturer’s protocol (10X User Guide). Generation of gel beads in emulsion (GEMs), barcoding, GEM-RT clean-up, complementary DNA amplification and library construction were all performed as per the manufacturer’s protocol. Individual sample quality was checked using a Bioanalyzer Tapestation (Agilent). Qubit was used for library quantification before pooling. The final library pool was sequenced on the Illumina HiSeq 4000 instrument using 50bp paired-end reads.

### Quantification and statistical analysis

#### General

Desmosome separation measurements in uninfected, distant to worm and adjacent (infected) cells in caecaloids and percentages of *Isg15^+^* crypts in uninfected and infected mice after 24 and 72h p.i. were compared using Kruskal Wallis and Dunn’s comparison tests from the Prism 8.2 software (GraphPad). Statistical comparison for desmosome separation in uninfected and infected mice and the *in vitro* and *in vivo* models was done using Mann-Whitney tests from the Prism 8.2 software (GraphPad).

#### L1 and L2 larvae RNA-seq data processing and analysis

*T. muris* reference genome (PRJEB126) was downloaded from Wormbase Parasite v14^60^. Reads from each RNA-seq sample were mapped against predicted transcripts using Kallisto v0.42.3^61^. Indexing and quantification were performed using default parameters. Read counts from multiple transcripts were combined at the gene level. Genes with zero counts across all samples and samples with less than 500000 read counts were removed prior to analysis. Differentially expressed genes were determined using DESeq2 v1.22.2^62^. Contrasts were performed between egg samples and 3h L1s, 3h L1s and 24h L1s, and 24h L1s and L2s. Genes were identified as differentially expressed with an adjusted p-value of < 0.05 and a log2 fold change > 1 or < −1. To identify functional patterns in an unbiased way, GO terms were determined which were enriched in differentially expressed genes using TopGO v2.34.0^63^ (node size = 5, method = weight01, FDR = 0.05, statistic = Fisher).

Gene expression heatmaps were created by combining read counts across biological replicates and calculating log2(FPKM + 1) for each gene, in each condition. These data were plotted using pheatmap v1.0.11 in R for genes which were differentially expressed in any contrast and had the GO term GO:0004252 (serine-type endopeptidase activity), contained a WAP domain (Pfam: PF00095), a Kunitz domain (Pfam: PF00014) or a serpin domain (Pfam: PF00079).

#### Caecum and IEC bulk RNA-seq data processing and analysis

For bulk RNA-seq data from caecum and IEC samples, reads were pseudo-aligned to the mouse transcriptome (Ensembl release 98) using Kallisto v0.46.2^61^. The DESeq2 package (v1.26.0)^62^ was used to identify differentially expressed genes over the time course (caecum; likelihood ratio test) or by pairwise comparison with time-matched controls (IECs; Wald test). All differentially expressed genes with an FDR adjusted p-value < 0.05 are reported in *Supplementary Data 1* (caecum) and *2* (IECs). Significantly enriched GO terms (Biological Process) annotated to the differentially expressed genes were identified using the GOseq package (v1.38.0)^64^, accounting for gene length bias, and p-values FDR-corrected with the Benjamini-Hochberg method. Expressed genes were ranked by log(pvalue) (most significantly upregulated to most significantly downregulated) and the fgsea package (v1.12.0)^65^ used for pathway enrichment analysis with the MSigDB *M. musculus* Hallmark pathways^66^.

### 10x single cell RNA-seq analysis

#### Quality Control

Raw unfiltered count matrices were generated using the CellRanger software suite (v3.0.2), aligning against the mm10-3.0.0 (Ensembl 93) mouse reference transcriptome. Unless otherwise stated, all further analysis was performed using the Seurat package (v3.0.2)^67^ in R (v3.6.1). Each library was examined for its distribution of unique molecular identifiers (UMIs) per cell *(Supplementary Fig. 7a).* UMI filtering thresholds were set for each library at the first local minimum on a UMI density plot, with GEMs associated with fewer UMIs being excluded from further analysis. Cells with a high percentage (>30%) of UMIs associated with mitochondrial genes were also excluded *(Supplementary Figs. 7b* and *c).* All libraries were initially merged and clustered prior to doublet detection with the DoubletDecon package (v1.1.4)^68^. The Main_Doublet_Decon function was run 10 times, and the intersection of identified cells were considered to be doublets and removed from the dataset. A distinct cluster of contaminating immune cells (identified by expression of CD3 and CD45) was also removed. Post QC, average cell recovery was 1401 cells per sample, with a total of 22422 cells captured at a mean depth of 39336 UMIs per cell and 4910 mean genes per cell.

#### Clustering, cell type identification and pseudotime analysis

Libraries derived from all mice (four per condition) were merged. Feature counts were log-normalized and scaled, and the 2000 most highly variable genes (identified with the FindVariableFeatures function) used for dimensionality reduction with PCA. The first 35 principal components were used for running UMAP for visualisation (min.dist=0.05) and for neighbour finding, and clusters initially identified with the FindClusters function (resolution=0.6). We confirmed the absence of batch effects visually (*Supplementary Fig. Fig. 7d)*. Cluster specific marker genes were identified based on cells derived from the eight control mice using the FindAllMarkers function (logfc.threshold=0.5, only.pos=TRUE. min.pct=0.3,test.use=”wilcox”). Clusters were inspected and some very similar clusters were merged, leaving 11 distinct clusters, and cluster markers recalculated. Cluster sizes were quantified as a proportion of the total number of cells per mouse, and differences between control and infected (timepoint-matched) mice assessed with a two-tailed t test. Significantly enriched GO terms associated with cluster marker genes *(Supplementary Data 3)* were identified with the enrichGO function from the clusterProfiler package (v3.14.0).

To scrutinise the undifferentiated cells further, three undifferentiated clusters were isolated *in silico*. Feature counts were normalized and scaled, top 2000 variable features identified and PCA performed. The first 30 principal components were used for running UMAP for visualisation (min.dist=0.05) and for neighbour finding, and clusters identified with the FindClusters function (resolution=0.4). After merging of very similar clusters, five clusters remained and markers were recalculated as described above.

Libraries from eight control mice were further analysed with the monocle3 (v0.2.1) package^69^ to place the cells in pseudotime. We applied a standard workflow with preprocess_cds (num_dim=100), align_cds (correcting for batch), reduce_dimension (reduction_method=UMAP), cluster_cells and learn_graph to fit a trajectory graph. Two main partitions were identified, consisting of undifferentiated cells/enterocytes and goblet cells. Cells in the G2M phase were selected as the root of the trajectories for pseudotime ordering. We used partition-based graph abstraction (PAGA)^70^, implemented in SCANPY^71^, to draw an abstract connectivity graph. After standard pre-processing, regressing out sample batches, the PAGA graph was computed and visualized (threshold=0.3).

## Data availability

The transcriptomic datasets generated during and analysed the current study are available in the European Nucleotide Archive (ENA) repository (https://www.ebi.ac.uk/ena/browser/home), under the accession numbers ERP008531 (STDY 3371 - Mouse and *Trichuris muris* transcriptome time course), ERP126662 (STDY 4023 - Investigating the transcriptome of early infective larvae stage of *Trichuris muris*) and ERP021944 (STDY 4672 - Investigating early host intestinal epithelial cells - whipworm interactions).

## Acknowledgements

This work was supported by the National Centre for the Replacement, Refinement and Reduction of Animals in Research (UK) David Sainsbury Fellowship Grant NC/P001521/1; the Medical Research Council Grant MR/R002800/1; and Wellcome via core funding to the Wellcome Sanger Institute (grant 206194) and the Wellcome Centre for Cell-Matrix Research, University of Manchester (grant 088785/Z/09/Z).

## Author contributions

Conceptualization, M.A.D-C., R.K.G. and M.B.; data curation, M.A.D-C., D.G., F.H.R., K.R., J.A.G., A.J.R., C.S., C.R., and T.S.; formal analysis, M.A.D-C., D.G., F.H.R., K.R., J.A.G., A.J.R., C.S., and C.R.; funding acquisition, M.A. D-C., D.J.T., R.K.G and M.B.; investigation, M.A.D-C., D.G., C.C., K.R., J.A.G., A.J.B., H.M.B., M.L., A.S., P.S., N.R., C.M., C.B., C.S., C.R., J.G. M., C.M. C., T.S., and K.S.H.; methodology, M.A.D-C., D.G., F.H.R., K.R., J.A.G., A.J.B., H.M.B., A.J.R., A.S, P.S., N.R., C.S., C.R., T.S., and K.S.H.; project administration, M.A. D-C., N. H., M.S., and M.B.; resources, M.A. D-C., J.A.G., D.J.T., R.K.G and M.B.; supervision, M.A.D-C., D.J.T., R.K.G and M. B; validation, M.A.D-C., D.G., K.R., J.A.G., C.S., and C.R.; visualization, M.A.D-C., D.G., F.H.R., K.R., J.A.G., A.J.R., C.S., and C.R.; writing of the original draft, M.A.D-C., F.H.R., K.R., J.A.G., H.M.B., A.J.R., D.J.T., R.K.G. and M.B.; writing–review and editing, M.A.D-C., F.H.R., D.J.T., R.K.G. and M.B.

## Competing interests

The authors declare no competing interests.

## Notes

### Competing Interest Statement

The authors have declared no competing interest.

### Summary of Updates

Addition of new results, both in text and figures

## References

1. Else, K. J. et al. Whipworm and roundworm infections. Nat Rev Dis Primers 6, 44, doi:10.1038/s41572-020-0171-3 (2020).

2. Jourdan, P. M., Lamberton, P. H. L., Fenwick, A. & Addiss, D. G. Soil-transmitted helminth infections. Lancet 391, 252–265, doi:10.1016/S0140-6736(17)31930-X (2018).

3. Klementowicz, J. E., Travis, M. A. & Grencis, R. K. Trichuris muris: a model of gastrointestinal parasite infection. Seminars in immunopathology 34, 815–828, doi:10.1007/s00281-012-0348-2 (2012).

4. Panesar, T. S. & Croll, N. A. The location of parasites within their hosts: site selection by Trichuris muris in the laboratory mouse. Int J Parasitol 10, 261–273, doi:10.1016/0020-7519(80)90006-5 (1980).

5. Tilney, L. G., Connelly, P. S., Guild, G. M., Vranich, K. A. & Artis, D. Adaptation of a nematode parasite to living within the mammalian epithelium. J Exp Zool A Comp Exp Biol 303, 927–945, doi:10.1002/jez.a.214 (2005).

6. Hayes, K. S. et al. Exploitation of the intestinal microflora by the parasitic nematode Trichuris muris. Science 328, 1391–1394, doi:10.1126/science.1187703 (2010).

7. Wakelin, D. The development of the early larval stages of Trichuris muris in the albino laboratory mouse. J Helminthol 43, 427–436, doi:10.1017/s0022149x00004995 (1969).

8. Panesar, T. S. The early phase of tissue invasion by Trichuris muris (nematoda: Trichuroidea). Z Parasitenkd 66, 163–166, doi:10.1007/bf00925723 (1981).

9. Beer, R. J. Studies on the biology of the life-cycle of Trichuris suis Schrank, 1788. Parasitology 67, 253-262, doi:10.1017/s0031182000046497 (1973).

10. Gehart, H. & Clevers, H. Tales from the crypt: new insights into intestinal stem cells. Nat Rev Gastroenterol Hepatol 16, 19–34, doi:10.1038/s41575-018-0081-y (2019).

11. Lee, T. D. & Wright, K. A. The morphology of the attachment and probable feeding site of the nematode Trichuris muris (Schrank, 1788) Hall, 1916. Can J Zool 56, 1889-1905, doi:10.1139/z78-258 (1978).

12. O’Sullivan, J. D. B., Cruickshank, S. M., Starborg, T., Withers, P. J. & Else, K. J. Characterisation of cuticular inflation development and ultrastructure in Trichuris muris using correlative X-ray computed tomography and electron microscopy. Sci Rep 10, 5846, doi:10.1038/s41598-020-61916-0 (2020).

13. Grencis, R. K. Immunity to helminths: resistance, regulation, and susceptibility to gastrointestinal nematodes. Annu Rev Immunol 33, 201–225, doi:10.1146/annurev-immunol-032713-120218 (2015).

14. Artis, D. & Grencis, R. K. The intestinal epithelium: sensors to effectors in nematode infection. Mucosal immunology 1, 252–264, doi:10.1038/mi.2008.21 (2008).

15. Iglesias-Guimarais, V. et al. IFN-Stimulated Gene 15 Is an Alarmin that Boosts the CTL Response via an Innate, NK Cell-Dependent Route. J Immunol 204, 2110–2121, doi:10.4049/jimmunol.1901410 (2020).

16. Perng, Y. C. & Lenschow, D. J. ISG15 in antiviral immunity and beyond. Nat Rev Microbiol 16, 423–439, doi:10.1038/s41579-018-0020-5 (2018).

17. Ostvik, A. E. et al. Intestinal epithelial cells express immunomodulatory ISG15 during active ulcerative colitis and Crohn’s disease. J Crohns Colitis, doi:10.1093/ecco-jcc/jjaa022 (2020).

18. Duque-Correa, M. A. et al. Development of caecaloids to study host-pathogen interactions: new insights into immunoregulatory functions of Trichuris muris extracellular vesicles in the caecum. Int J Parasitol 50, 707–718, doi:10.1016/j.ijpara.2020.06.001 (2020).

19. McGuckin, M. A., Linden, S. K., Sutton, P. & Florin, T. H. Mucin dynamics and enteric pathogens. Nat Rev Microbiol 9, 265–278, doi:10.1038/nrmicro2538 (2011).

20. Hasnain, S. Z., McGuckin, M. A., Grencis, R. K. & Thornton, D. J. Serine protease(s) secreted by the nematode Trichuris muris degrade the mucus barrier. PLoS neglected tropical diseases 6, e1856, doi:10.1371/journal.pntd.0001856 (2012).

21. Lidell, M. E., Moncada, D. M., Chadee, K. & Hansson, G. C. Entamoeba histolytica cysteine proteases cleave the MUC2 mucin in its C-terminal domain and dissolve the protective colonic mucus gel. Proc Natl Acad Sci U S A 103, 9298–9303, doi:10.1073/pnas.0600623103 (2006).

22. Buckley, A. & Turner, J. R. Cell Biology of Tight Junction Barrier Regulation and Mucosal Disease. Cold Spring Harb Perspect Biol 10, doi:10.1101/cshperspect.a029314 (2018).

23. Haber, A. L. et al. A single-cell survey of the small intestinal epithelium. Nature 551, 333–339, doi:10.1038/nature24489 (2017).

24. Yan, K. S. et al. Intestinal Enteroendocrine Lineage Cells Possess Homeostatic and Injury-Inducible Stem Cell Activity. Cell Stem Cell 21, 78–90 e76, doi:10.1016/j.stem.2017.06.014 (2017).

25. Parikh, K. et al. Colonic epithelial cell diversity in health and inflammatory bowel disease. Nature 567, 49–55, doi:10.1038/s41586-019-0992-y (2019).

26. Murata, K. et al. Ascl2-Dependent Cell Dedifferentiation Drives Regeneration of Ablated Intestinal Stem Cells. Cell Stem Cell 26, 377–390 e376, doi:10.1016/j.stem.2019.12.011 (2020).

27. Wang, Y. et al. Single-cell transcriptome analysis reveals differential nutrient absorption functions in human intestine. J Exp Med 217, doi:10.1084/jem.20191130 (2020).

28. Herring, C. A. et al. Unsupervised Trajectory Analysis of Single-Cell RNA-Seq and Imaging Data Reveals Alternative Tuft Cell Origins in the Gut. Cell Syst 6, 37–51 e39, doi:10.1016/j.cels.2017.10.012 (2018).

29. Liu, Q. et al. Quantitative assessment of cell population diversity in single-cell landscapes. PLoS Biol 16, e2006687, doi:10.1371/journal.pbio.2006687 (2018).

30. Bancroft, A. J. et al. The major secreted protein of the whipworm parasite tethers to matrix and inhibits interleukin-13 function. Nat Commun 10, 2344, doi:10.1038/s41467-019-09996-z (2019).

31. Sudha, P. S., Devaraj, H. & Devaraj, N. Adherence of Shigella dysenteriae 1 to human colonic mucin. Curr Microbiol 42, 381–387, doi:10.1007/s002840010234 (2001).

32. Ravdin, J. I. & Guerrant, R. L. Role of adherence in cytopathogenic mechanisms of Entamoeba histolytica. Study with mammalian tissue culture cells and human erythrocytes. J Clin Invest 68, 1305–1313, doi:10.1172/jci110377 (1981).

33. Chadee, K., Petri, W. A., Jr., Innes, D. J. & Ravdin, J. I. Rat and human colonic mucins bind to and inhibit adherence lectin of Entamoeba histolytica. J Clin Invest 80, 1245–1254, doi:10.1172/JCI113199 (1987).

34. Hasnain, S. Z. et al. Immune-driven alterations in mucin sulphation is an important mediator of Trichuris muris helminth expulsion. PLoS Pathog 13, e1006218, doi:10.1371/journal.ppat.1006218 (2017).

35. Tort, J., Brindley, P. J., Knox, D., Wolfe, K. H. & Dalton, J. P. Proteinases and associated genes of parasitic helminths. Advances in parasitology 43, 161–266, doi:10.1016/s0065-308x(08)60243-2 (1999).

36. Dzik, J. M. Molecules released by helminth parasites involved in host colonization. Acta Biochim Pol 53, 33–64 (2006).

37. Gause, W. C., Rothlin, C. & Loke, P. Heterogeneity in the initiation, development and function of type 2 immunity. Nature reviews. Immunology 20, 603–614, doi:10.1038/s41577-020-0301-x (2020).

38. Owyang, A. M. et al. Interleukin 25 regulates type 2 cytokine-dependent immunity and limits chronic inflammation in the gastrointestinal tract. J Exp Med 203, 843–849, doi:10.1084/jem.20051496 (2006).

39. Saenz, S. A. et al. IL25 elicits a multipotent progenitor cell population that promotes T(H)2 cytokine responses. Nature 464, 1362–1366, doi:10.1038/nature08901 (2010).

40. Humphreys, N. E., Xu, D., Hepworth, M. R., Liew, F. Y. & Grencis, R. K. IL-33, a potent inducer of adaptive immunity to intestinal nematodes. J Immunol 180, 2443–2449, doi:10.4049/jimmunol.180.4.2443 (2008).

41. Chen, Z. et al. Interleukin-33 Promotes Serotonin Release from Enterochromaffin Cells for Intestinal Homeostasis. Immunity, doi:10.1016/j.immuni.2020.10.014 (2020).

42. Taylor, B. C. et al. TSLP regulates intestinal immunity and inflammation in mouse models of helminth infection and colitis. J Exp Med 206, 655–667, doi:10.1084/jem.20081499 (2009).

43. Zaph, C. et al. Epithelial-cell-intrinsic IKK-beta expression regulates intestinal immune homeostasis. Nature 446, 552–556, doi:10.1038/nature05590 (2007).

44. Werneke, S. W. et al. ISG15 is critical in the control of Chikungunya virus infection independent of UbE1L mediated conjugation. PLoS Pathog 7, e1002322, doi:10.1371/journal.ppat.1002322 (2011).

45. Morales, D. J. et al. Novel mode of ISG15-mediated protection against influenza A virus and Sendai virus in mice. J Virol 89, 337–349, doi:10.1128/JVI.02110-14 (2015).

46. Heo, I. et al. Modelling Cryptosporidium infection in human small intestinal and lung organoids. Nat Microbiol 3, 814–823, doi:10.1038/s41564-018-0177-8 (2018).

47. Dong, C., Gao, N., Ross, B. X. & Yu, F. X. ISG15 in Host Defense Against Candida albicans Infection in a Mouse Model of Fungal Keratitis. Invest Ophthalmol Vis Sci 58, 2948–2958, doi:10.1167/iovs.17-21476 (2017).

48. Nikolaev, M. et al. Homeostatic mini-intestines through scaffold-guided organoid morphogenesis. Nature 585, 574–578, doi:10.1038/s41586-020-2724-8 (2020).

49. Radoshevich, L. et al. ISG15 counteracts Listeria monocytogenes infection. Elife 4, doi:10.7554/eLife.06848 (2015).

50. Webb, L. M. et al. Type I interferon is required for T helper (Th) 2 induction by dendritic cells. EMBO J 36, 2404–2418, doi:10.15252/embj.201695345 (2017).

51. Connor, L. M. et al. Th2 responses are primed by skin dendritic cells with distinct transcriptional profiles. J Exp Med 214, 125–142, doi:10.1084/jem.20160470 (2017).

52. Reynolds, L. A. et al. MyD88 signaling inhibits protective immunity to the gastrointestinal helminth parasite Heligmosomoides polygyrus. J Immunol 193, 2984–2993, doi:10.4049/jimmunol.1401056 (2014).

53. Wakelin, D. Acquired immunity to Trichuris muris in the albino laboratory mouse. Parasitology 57, 515–524 (1967).

54. Marconi, A., Hancock-Ronemus, A. & Gillis, J. A. Adult chondrogenesis and spontaneous cartilage repair in the skate, Leucoraja erinacea. Elife 9, doi:10.7554/eLife.53414 (2020).

55. Choi, H. M. T. et al. Third-generation in situ hybridization chain reaction: multiplexed, quantitative, sensitive, versatile, robust. Development 145, doi:10.1242/dev.165753 (2018).

56. Criswell, K. E. & Gillis, J. A. Resegmentation is an ancestral feature of the gnathostome vertebral skeleton. Elife 9, doi:10.7554/eLife.51696 (2020).

57. Malick, L. E. & Wilson, R. B. Modified thiocarbohydrazide procedure for scanning electron microscopy: routine use for normal, pathological, or experimental tissues. Stain Technol 50, 265–269, doi:10.3109/10520297509117069 (1975).

58. Deerinck, T. J., Bushong, E. A., Lev-Ram, V., Shu, X., Tsien, R.Y. and Ellisman, M. H.. Enhancing Serial Block-Face Scanning Electron Microscopy to Enable High Resolution 3-D Nanohistology of Cells and Tissues. Microscopy and Microanalysis 16 1138-1139, doi:doi:10.1017/S1431927610055170 (2010).

59. Picelli, S. et al. Full-length RNA-seq from single cells using Smart-seq2. Nat Protoc 9, 171–181, doi:10.1038/nprot.2014.006 (2014).

60. Bolt, B. J. et al. Using WormBase ParaSite: An Integrated Platform for Exploring Helminth Genomic Data. Methods Mol Biol 1757, 471–491, doi:10.1007/978-1-4939-7737-6_15 (2018).

61. Bray, N. L., Pimentel, H., Melsted, P. & Pachter, L. Near-optimal probabilistic RNA-seq quantification. Nat Biotechnol 34, 525–527, doi:10.1038/nbt.3519 (2016).

62. Love, M. I., Huber, W. & Anders, S. Moderated estimation of fold change and dispersion for RNA-seq data with DESeq2. Genome Biol 15, 550, doi:10.1186/s13059-014-0550-8 (2014).

63. Alexa, A., Rahnenfuhrer, J. & Lengauer, T. Improved scoring of functional groups from gene expression data by decorrelating GO graph structure. Bioinformatics 22, 1600–1607, doi:10.1093/bioinformatics/btl140 (2006).

64. Young, M. D., Wakefield, M. J., Smyth, G. K. & Oshlack, A. Gene ontology analysis for RNA-seq: accounting for selection bias. Genome Biol 11, R14, doi:10.1186/gb-2010-11-2-r14 (2010).

65. Korotkevich, G. et al. Fast gene set enrichment analysis. bioRxiv, 060012, doi:10.1101/060012 (2021).

66. Liberzon, A. et al. The Molecular Signatures Database (MSigDB) hallmark gene set collection. Cell Syst 1, 417–425, doi:10.1016/j.cels.2015.12.004 (2015).

67. Stuart, T. et al. Comprehensive Integration of Single-Cell Data. Cell 177, 1888–1902 e1821, doi:10.1016/j.cell.2019.05.031 (2019).

68. DePasquale, E. A. K. et al. DoubletDecon: Deconvoluting Doublets from Single-Cell RNA-Sequencing Data. Cell Rep 29, 1718–1727 e1718, doi:10.1016/j.celrep.2019.09.082 (2019).

69. Trapnell, C. et al. The dynamics and regulators of cell fate decisions are revealed by pseudotemporal ordering of single cells. Nat Biotechnol 32, 381–386, doi:10.1038/nbt.2859 (2014).

70. Wolf, F. A. et al. PAGA: graph abstraction reconciles clustering with trajectory inference through a topology preserving map of single cells. Genome Biol 20, 59, doi:10.1186/s13059-019-1663-x (2019).

71. Wolf, F. A., Angerer, P. & Theis, F. J. SCANPY: large-scale single-cell gene expression data analysis. Genome Biol 19, 15, doi:10.1186/s13059-017-1382-0 (2018).

