## Supplementary materials for "Defining the early stages of intestinal colonisation by whipworms"

**SUPPLEMENTARY FIGURES**

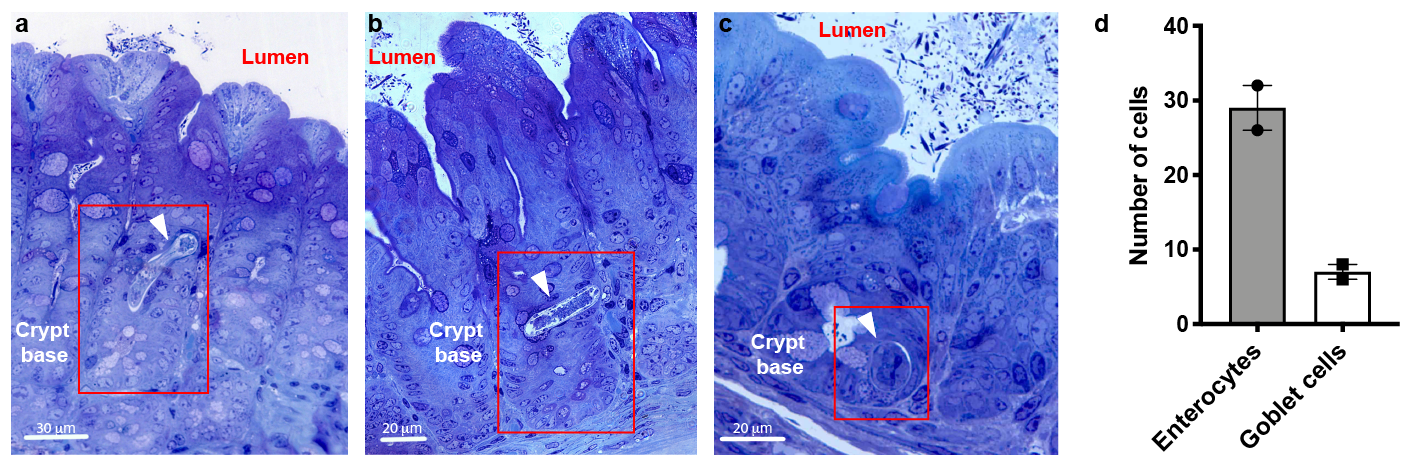

**Supplementary Fig. 1. Whipworm L1 larvae become completely intracellular in syncytial tunnels at the bottom of the crypts of Lieberkühn.**

**a-c** Images of toluidine blue-stained transverse sections from caecum of WT mice infected with *T. muris* at 3 h p.i., showing whipworm larvae (arrowhead) infecting cells at the base of crypts. **d** Numbers of enterocytes and goblet cells infected by whipworm L1 larvae in syncytial tunnels *in vivo* (24 h p.i.). Counts on serial block face SEM images of two syncytial tunnels. Points are the counts for each individual worm.

**Supplementary Fig. 2. Characterisation of caecaloids grown in transwells.**

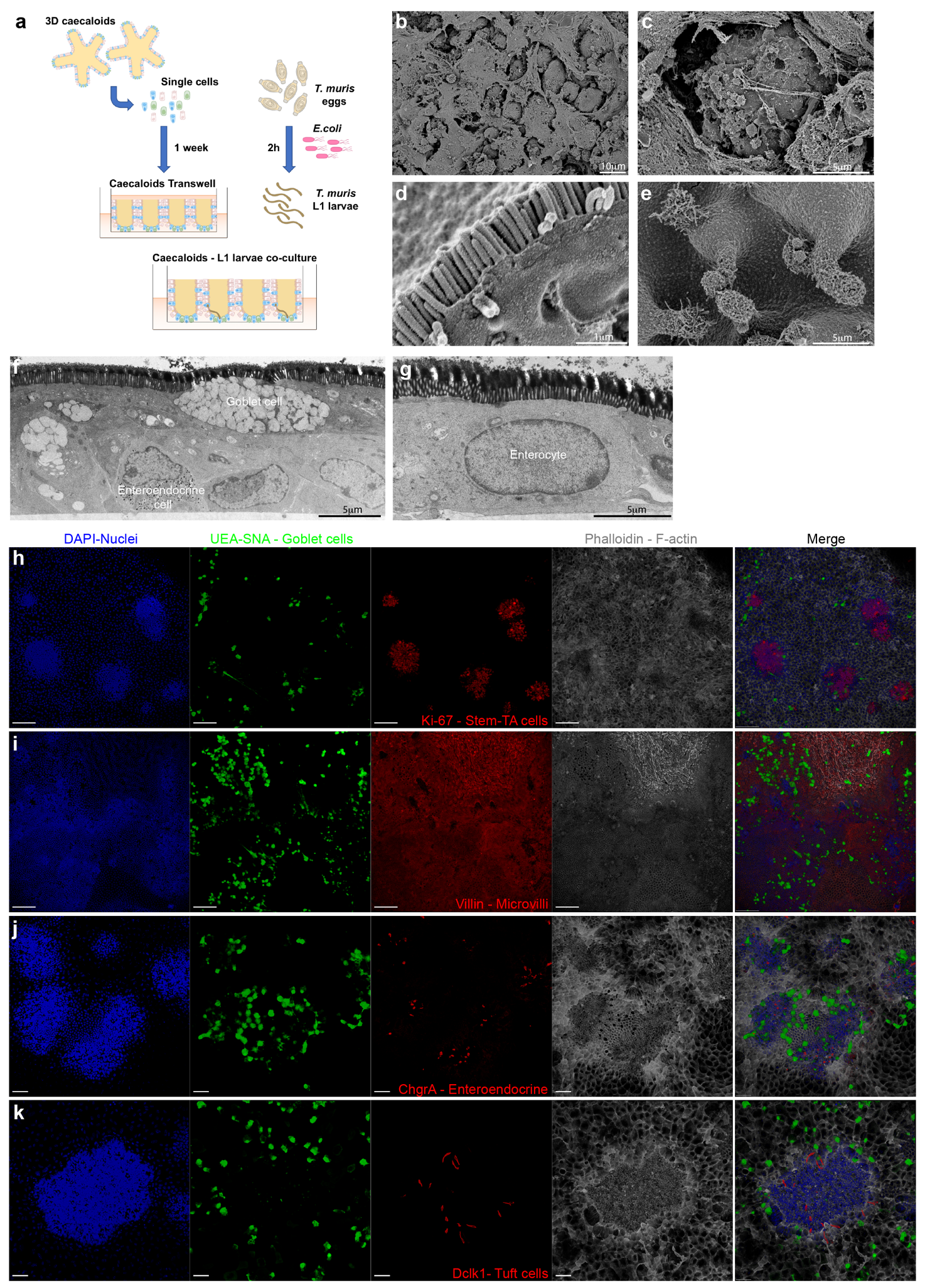

**a** Scheme of the caecaloid-whipworm model experimental set-up. **b-k** Images of caecaloids grown and differentiated in transwells showing the presence of all IEC populations organized in structures resembling intestinal crypts. **b-e** Scanning EM images showing **(b)** mucus and **(c)** goblet cells, **(d)** microvilli of enterocytes and **(e)** tuft cells. **f-g** Transmission EM showing **(f)** goblet and enteroendocrine cells and **(g)** enterocytes**. h-k** Confocal IF microscopy with antibodies staining **(h)** Ki-67, marker of proliferating cells, stem and TA cells; **(i)** villin, identifying microvilli of enterocytes; **(j)** chromogranin A expressing enteroendocrine cells; and **(k)** Dclk-1, marker of tuft cells. Lectins UEA and SNA bind mucin glycans in goblet cells. DAPI stains nuclei and Phalloidin binds to F-actin. Scale bar 100μm for h and i, 50μm for j and k.

**Supplementary Fig. 3. Expression of serine proteases and peptidase inhibitors by L1 larvae in early stages of infection.**

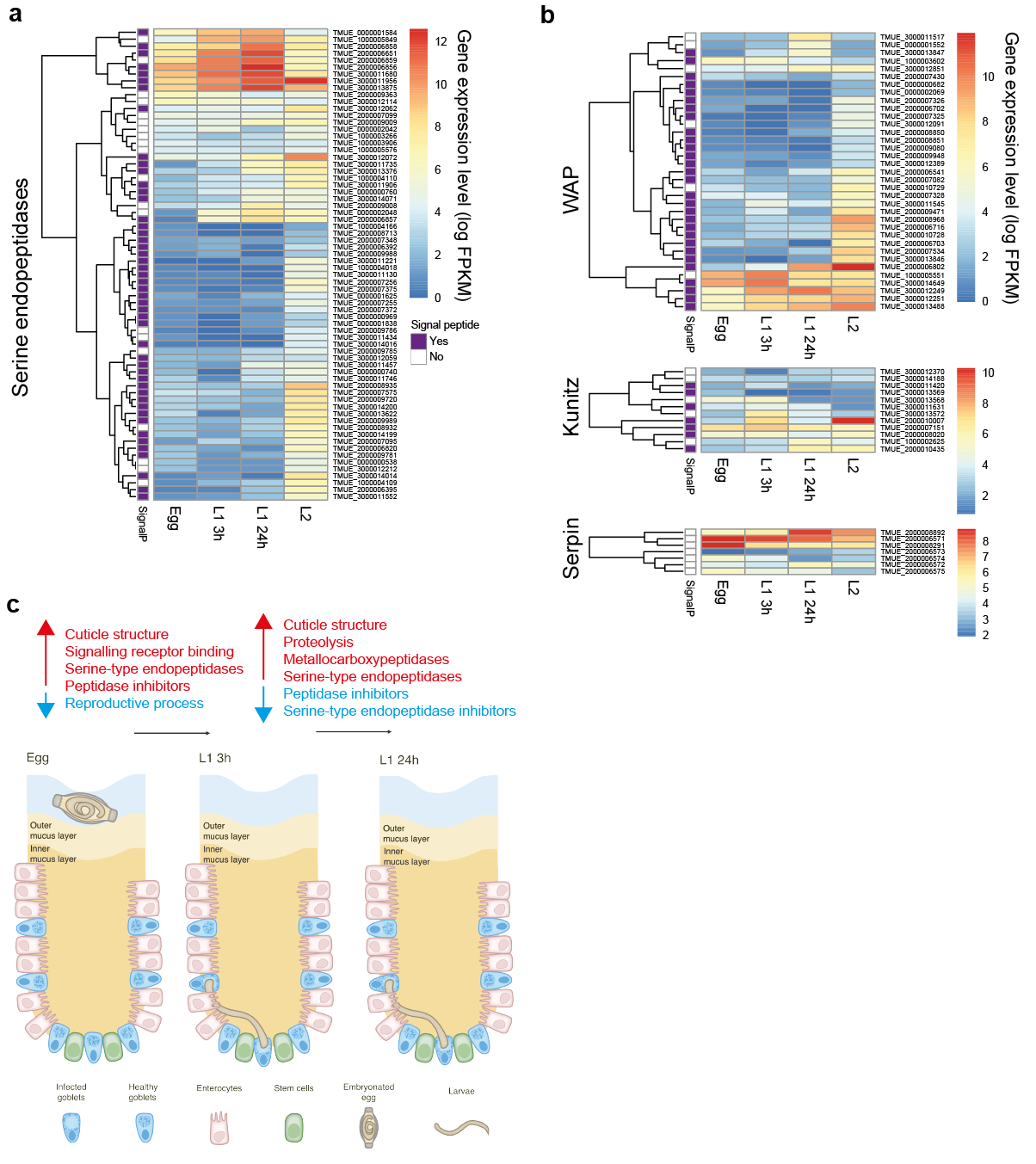

Expression levels of **a** serine endopeptidases and **b** WAP, Kunitz and serpin peptidase inhibitors from *T. muris* are shown over early larval development as log-normalised Fragments Per Kilobase per Million mapped reads (FPKM). Only genes differentially expressed across the time course are shown. **c** A diagram of early larval development in a caecal crypt is annotated with examples of GO terms enriched amongst genes with increasing transcript levels (red) or reduced transcript levels (blue) between stages.

**Supplementary Fig. 4. Mucus degradation by L1 larvae in early stages of infection.**

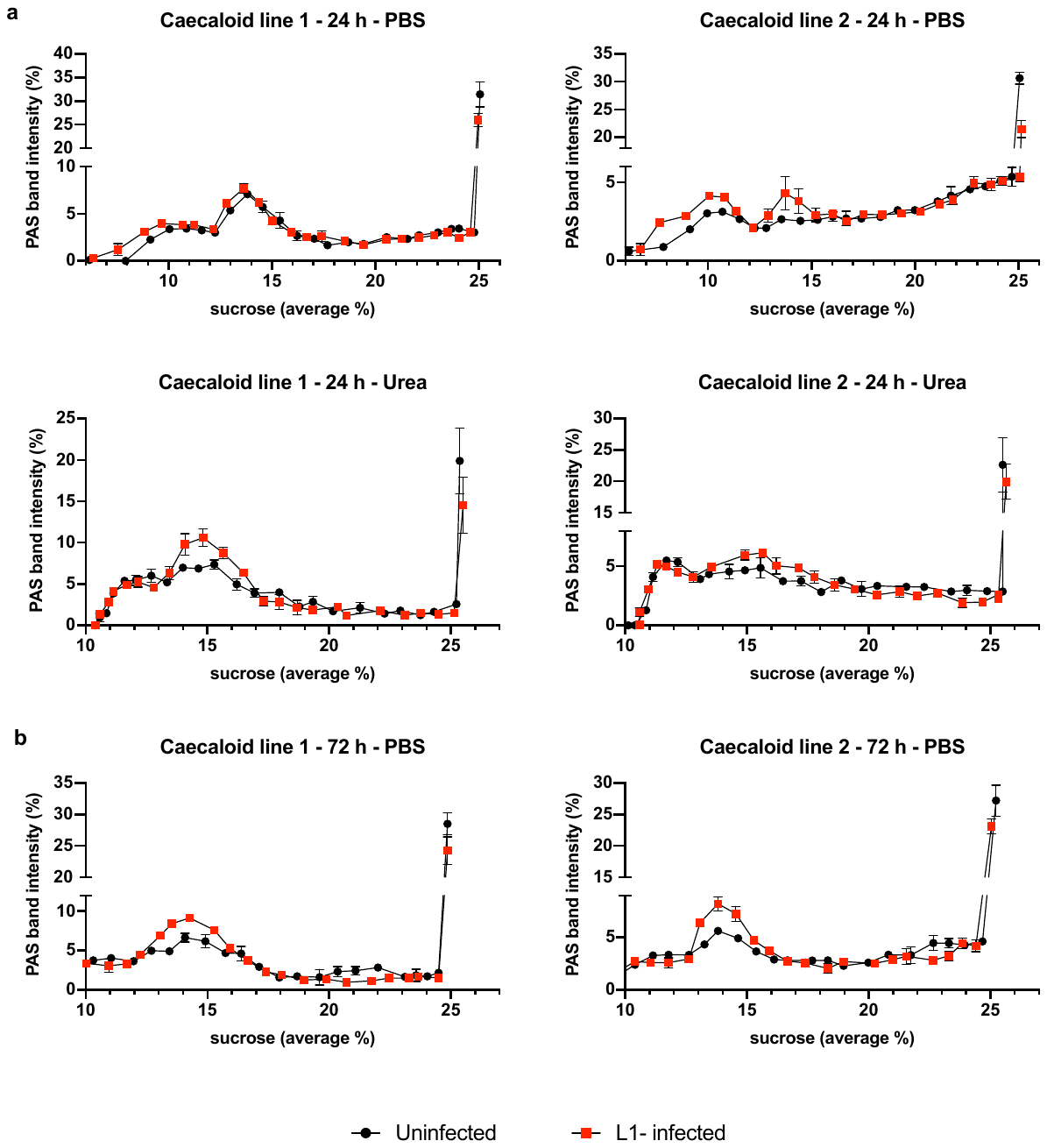

Caecaloid mucus degradation by L1 larvae at **a** 24 and **b** 72 h p.i. Transwells were washed with PBS followed by 0.2 M urea in PBS to recover mucus. Washes were subjected to rate zonal centrifugation on linear 5-25% sucrose gradients. After centrifugation tubes were emptied from the top and the fractions were stained with PAS to detect the mucins. Data are shown as percentage of intensity. Results are represented as the mean +/- SEM of 3 replicas of two caecaloid lines.

**Supplementary Fig. 5. Perturbation of host cell desmosomes, but not tight and adherens junctions, upon whipworm L1 larvae infection of mice and caecaloids.**

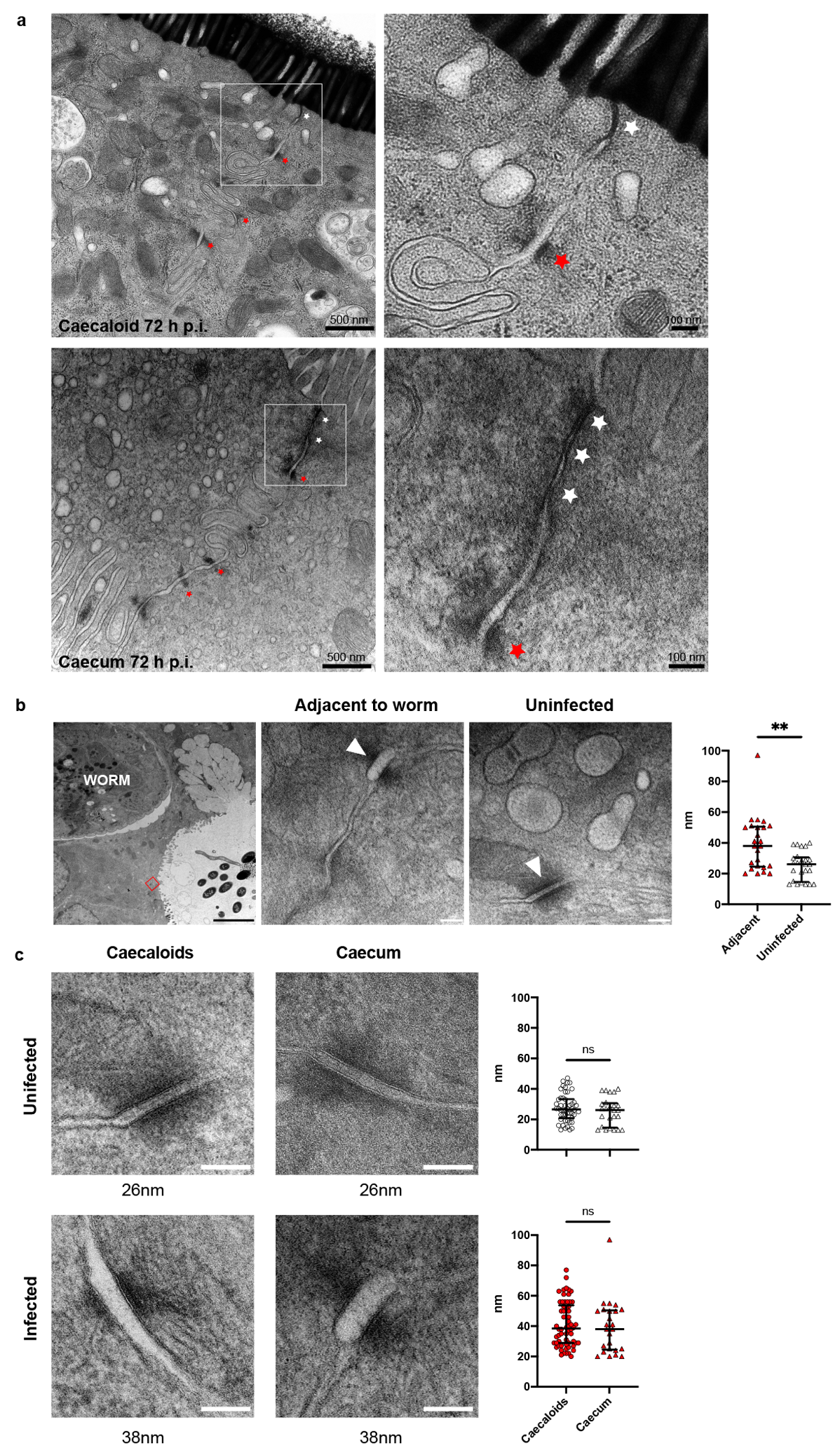

Representative TEM images of *T. muris-*infected caecaloids and caecum after 72 h of infection showing: **a** Intact tight and adherens junctions (white stars) but separated desmosomes (red stars) on cells hosting the larva; **b** Worm infecting IEC at the caecum (scale bar 3μm) and desmosomes (arrowheads) joining infected and adjacent cells from infected mice (inset) and cells from uninfected mice. Scale bars for desmosome images 100nm. Desmosome separation in nm was measured in host cells from two worms. Adjacent measurements n=25, uninfected measurements n=25. **p<0.0025 Mann-Whitney test. **c** Representative TEM images of desmosomes from uninfected and *T. muris-*infected caecaloids and caecum of mice. Scale bars 100nm. Statistical comparison between the desmosome separation measurements show no difference between the *in vitro* and *in vivo* models. Measurements: caecaloid uninfected n=50, caecum infected n=25, caecaloid infected n=62, caecum infected n=25. Mann-Whitney test.

**Supplementary Fig. 6. Host responses to early infection with whipworms are dominated by a type-I IFN signature.**

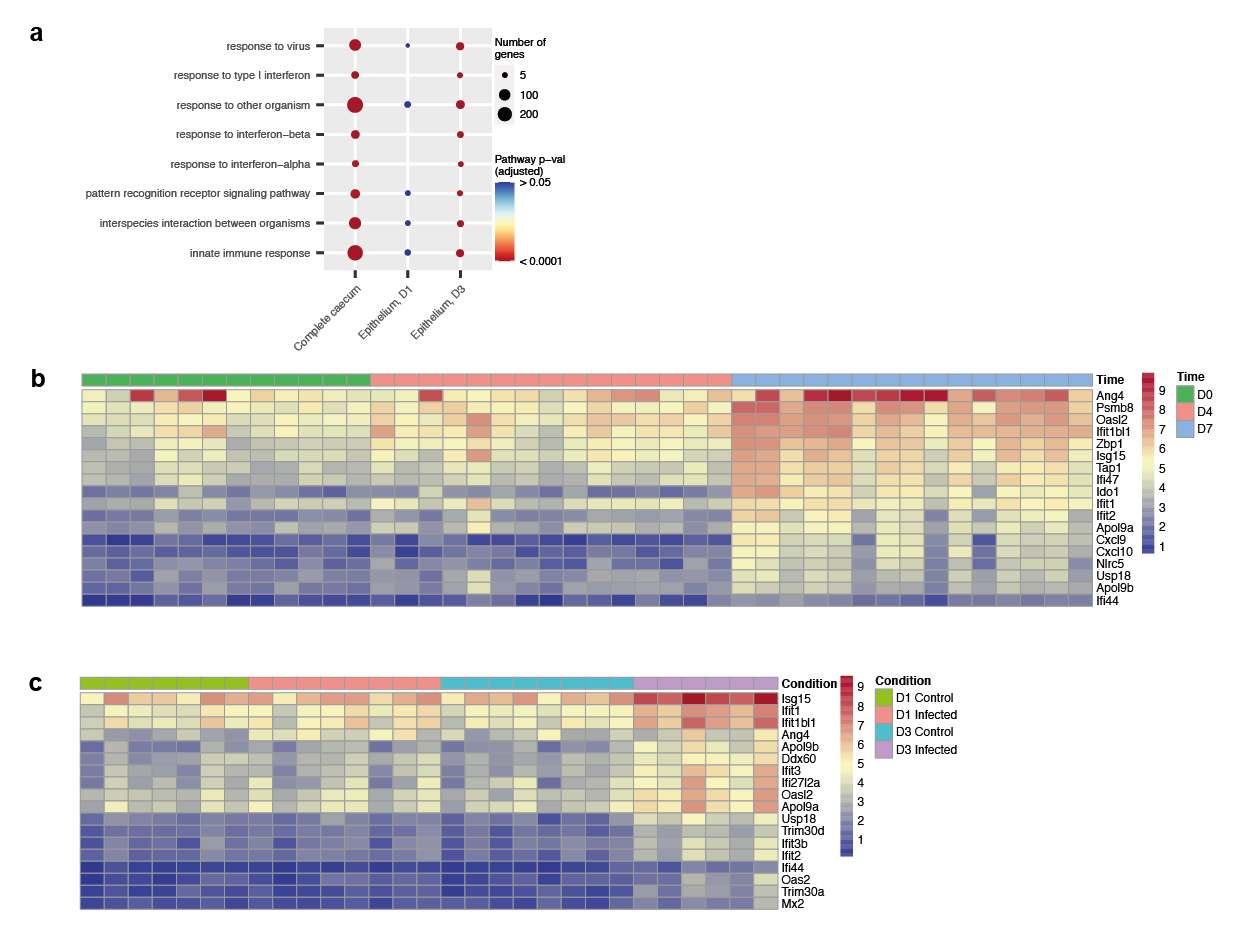

**a** Selected significantly enriched gene ontology (GO) terms associated with differentially expressed genes in the complete caecum (7 day *T. muris* infection time course) and caecal IECs (D1 and D3 post *T. muris* infection). The size of each circle corresponds to the numbers of genes annotated to a given term, while the colour indicates the FDR-adjusted p value (n= 4-5 mice per group). **b** Heatmap (log2(FPKM+1)) of selected IFN-associated differentially expressed genes (all with log2FoldChange > 1.5 at 0 v 7 d.p.i and FDR-adjusted p value <0.05) in the caecum of mice in response to *T. muris* infection at days 0 (control), 4 and 7 p.i (n=4-5 mice per group, with 3 caecum samples per mouse). **c** Heatmap (log2(FPKM+1)) of selected IFN-associated differentially expressed genes (all with log2FoldChange > 1.5 at 3 d.p.i and FDR-adjusted p value <0.05) in caecal IECs at 1 and 3 d.p.i, with time-matched controls (n=4 mice per group, with 1-2 technical replicates per mouse).

**Supplementary Fig 7. Single cell RNA-seq data QC.**

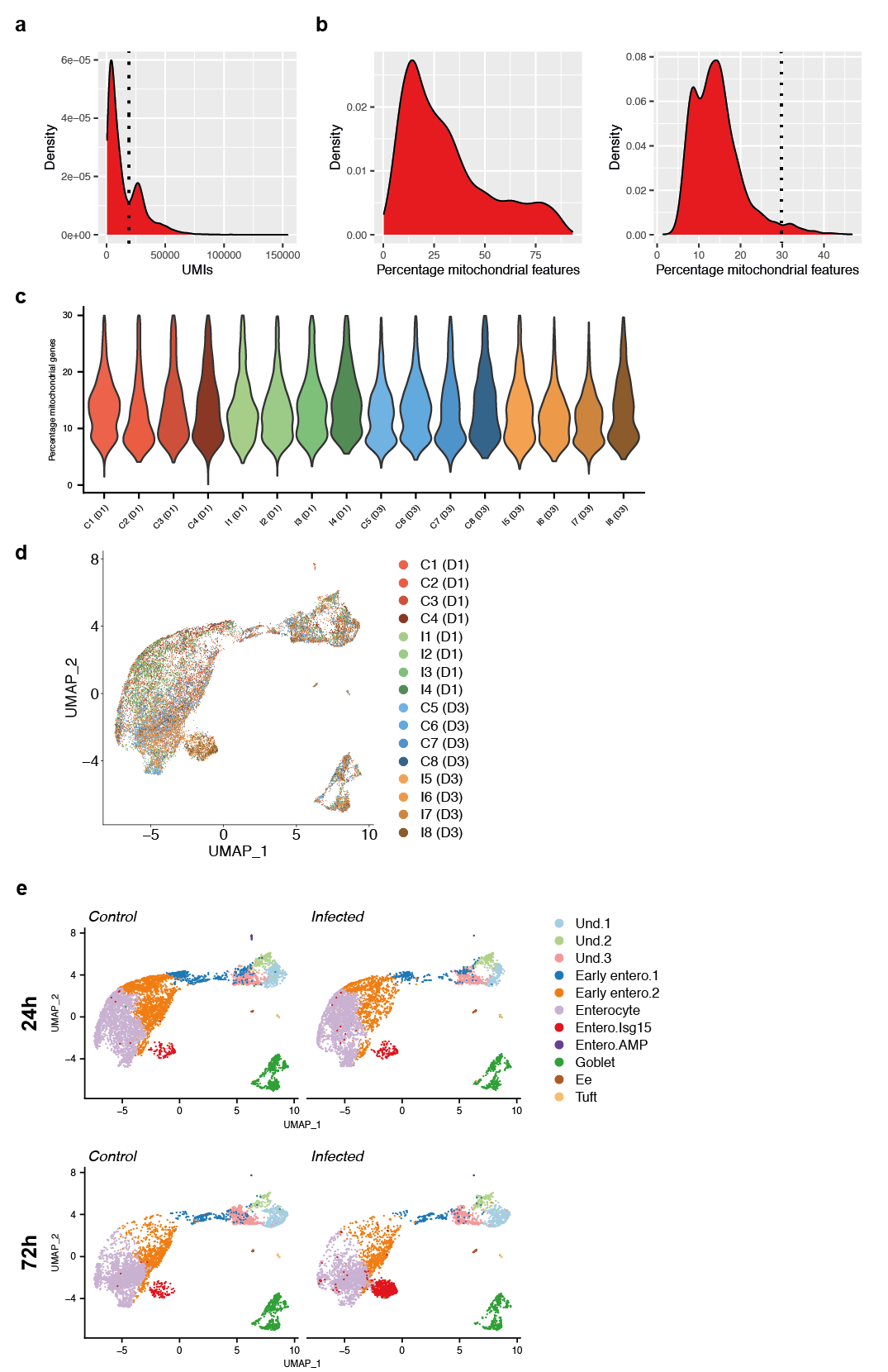

**a** Cellular barcodes in 10x data were selected above the first local minimum on a UMI density plot for each individual sample. Example distribution density and local minimum (dashed line) are shown. **b** Percentage of mitochondrial genes per cell for a representative sample pre (left panel) and post (right panel) UMI-based filtering. An additional filter at 30% mitochondrial genes per cell was then applied (dotted line). **c** Violin plots of percentage of mitochondrial genes per cell post QC (all samples; C= control, I=infected). **d** UMAP visualization of sample batch distribution in the merged clustering analysis (n = 4 mice per group). **e** UMAP plots from single cell RNA-seq analysis of IEC populations (colour coded) in the caecum of control and *T. muris*-infected mice after 1 and 3 days p.i. (n=4 mice for each condition at each time point).

**Supplementary Fig 8. *In silico* analysis of undifferentiated clusters.**

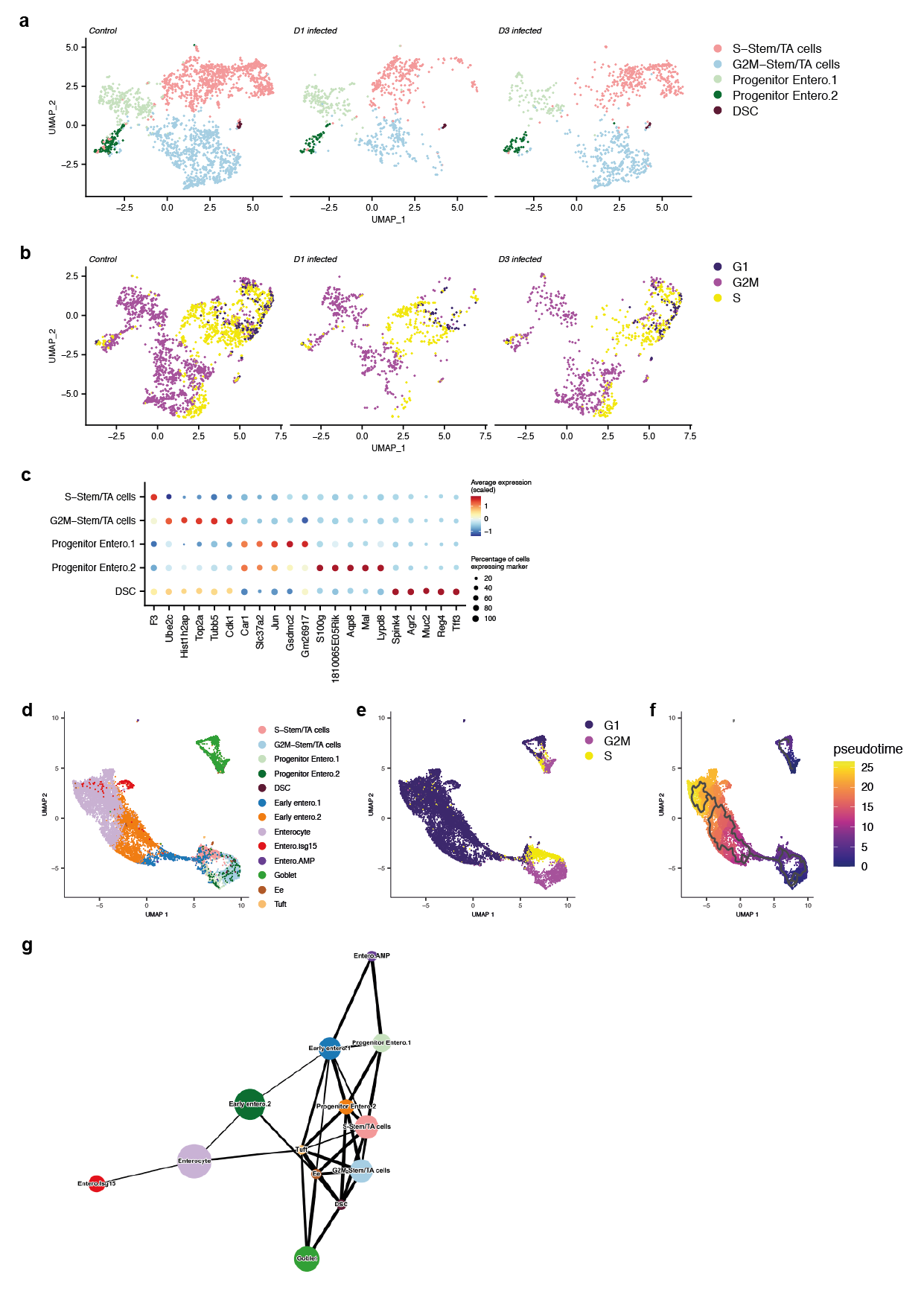

**a** UMAP visualisation of subclustered undifferentiated cells, coloured by subcluster. **b** UMAP visualisation of undifferentiated cells, coloured by cell cycle phase**. c** Dot plot of the top marker genes for each cell type. The relative size of each dot represents the fraction of cells per cluster that expresses each marker; the colour represents the average (scaled) gene expression per cluster. **d** Monocle UMAP visualisation of all control cells, coloured by cluster (integrating subclustering of undifferentiated cells). **e** Monocle UMAP visualisation of all control cells, coloured by cell cycle phase. **f** Monocle UMAP visualisation of all control cells, coloured by pseudotime. **g** Partition-based graph abstraction (PAGA) visualisation of all control cells. Line thickness represents relatedness between clusters.

**Supplementary Fig 9. Expression of stem, TA and progenitor cell markers by IECs from the caecum of uninfected mice.**

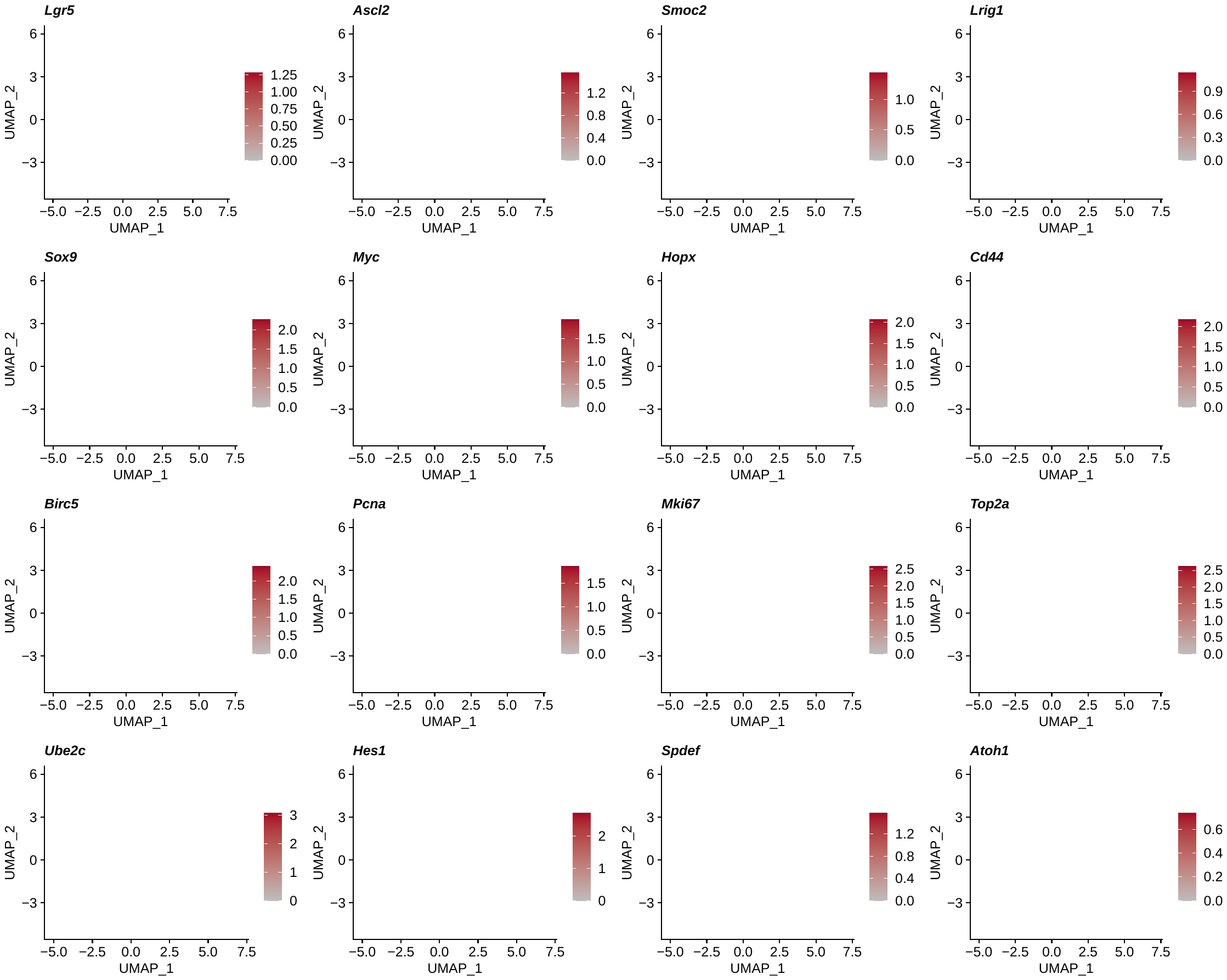

UMAP plots from single cell RNA-seq analysis showing normalized expression of *Lgr5*, *Ascl2,* *Smoc2, Lrig1*, *Sox9*, *Myc*, *Hopx, Cd44, Birc5, Pcna, Mki67, Top2a, Ubec2c, Hes1, Spdef* and *Atoh1* in IECs from the caecum of control mice (n=8).

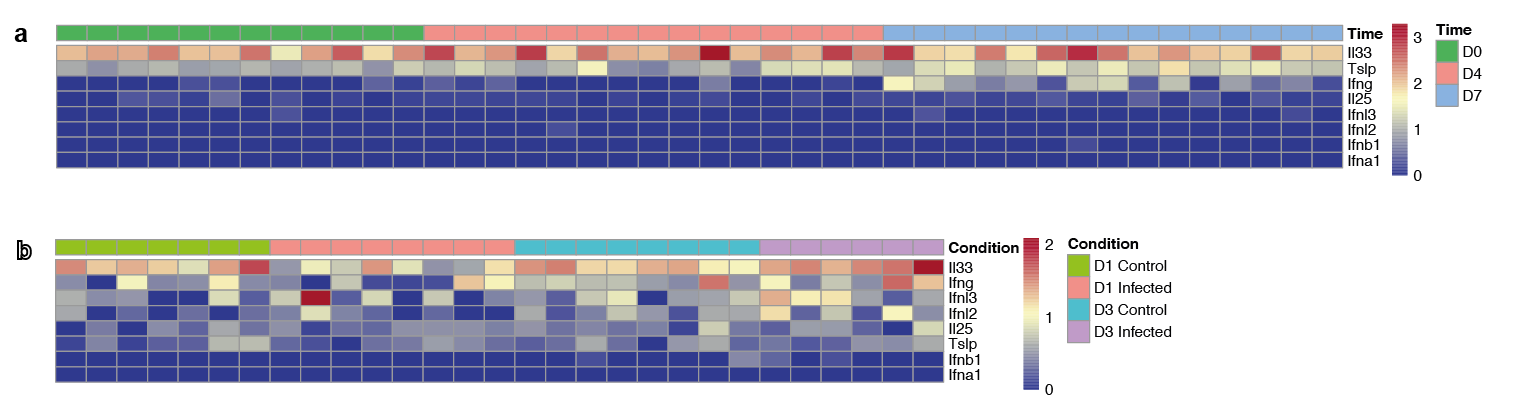

**Supplementary Fig 10. Expression of cytokines and alarmins in the caecum and caecal IECs of uninfected and *T. muris*-infected mice upon early infection.**

**a** Expression (log2(FPKM+1)) of *Il33*, *Tslp*, *Ifng,* *Il25*, *Ifnl3, Ifnl2, Ifnb1* and *Ifna1* in the caecum of mice at days 0, 4 and 7 post *T. muris* infection (n=4-5 mice per group, with 3 caecum samples per mouse). **b** Expression (log2(FPKM+1)) of *Il-33*, *Ifng, Ifnl3, Ifnl2, Il-25*, *Tslp, Ifnb1* and *Ifna1* in sorted murine caecal IECs in response to *T. muris* infection at days 1 and 3 p.i (n=4 mice per group, with 1-2 technical replicates per mouse).

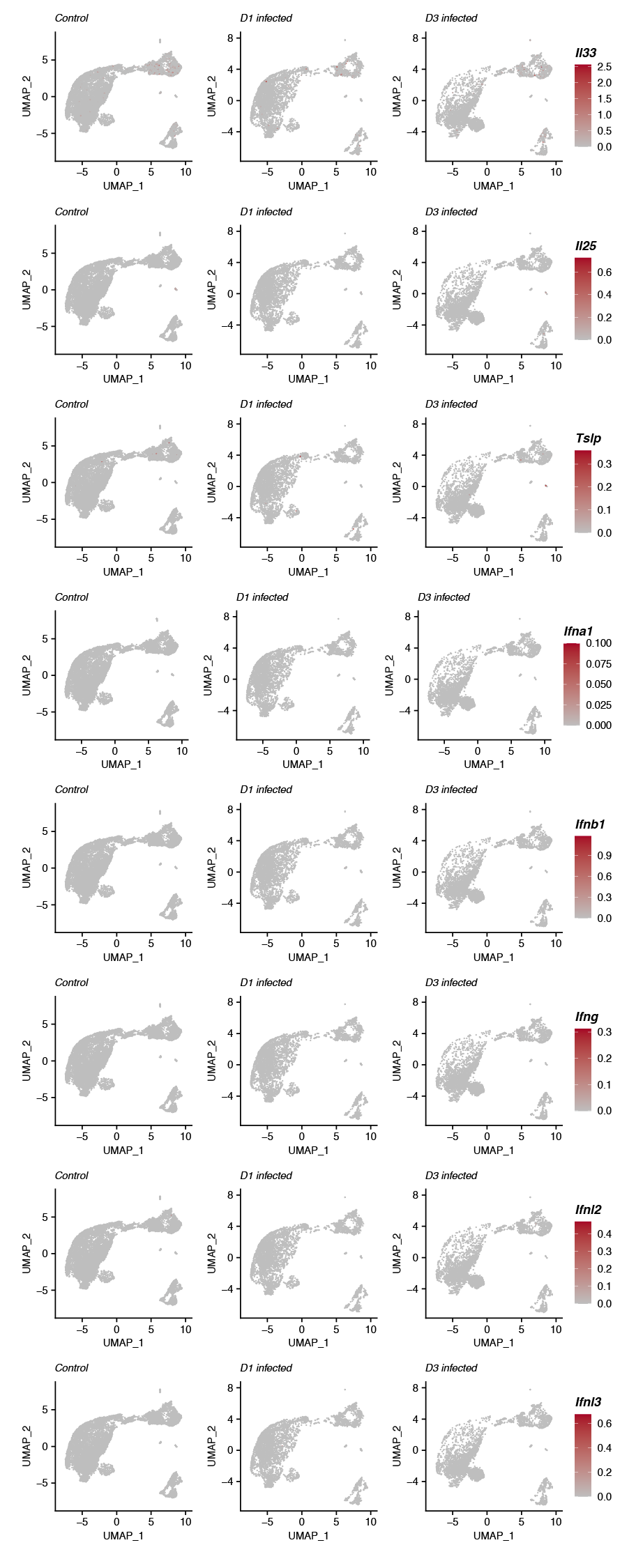
**Supplementary Fig 11. Expression of cytokines and alarmins by IECs from the caecum of uninfected and *T. muris*-infected mice upon early infection.**

UMAP plots from single cell RNA-seq analysis showing normalized expression levels of *Il33*, *Il25*, *Tslp*, *Ifna1*, *Ifnb1, Ifng,* *Ifnl2* and *Ifnl3* in IECs from the caecum of control (n=8, 4 mice for each time point) and *T. muris*-infected mice after 1 and 3 days p.i. (n=4 mice for each time point).

**SUPPLEMENTARY VIDEO LEGENDS**

**Supplementary videos 1 and 2. Visualisation of syncytial tunnels burrowed by *T. muris* L1 larvae infecting IECs *in vivo*.** Videos visualising serial block face SEM images of syncytial tunnels in the caecum of *T. muris*-infected mice at 24 h p.i.

**Supplementary Video 3. Visualisation of syncytial tunnel burrowed by *T. muris* L1 larva infecting IECs in caecaloids.** Video visualising z-stack of confocal IF images of caecaloids infected with whipworm L1 larvae for 24 h. Note the intricate tunnel left behind by larva, which are completely multi-intracellular. Stained in blue are nuclei of both the epithelia cells and larvae (DAPI), in white is F-actin at cell membrane (phalloidin), in green are mucus vacuoles of goblet cells (UEA/SNA lectins binding mucin glycans) and in magenta are dividing cells (Ki-67).

**Supplementary Video 4. Visualisation of syncytial tunnel burrowed by *T. muris* L1 larva infecting IECs in caecaloids.** Video visualising z-stack of confocal IF images of caecaloids infected with whipworm L1 larvae for 24 h. Note the intricate tunnel left behind by larva, which are completely multi-intracellular. Stained in blue are nuclei of both the epithelia cells and larvae (DAPI), in white is F-actin at cell membrane (phalloidin), in green are mucus vacuoles of goblet cells (UEA/SNA lectins binding mucin glycans) and in red are tuft cells (Dckl-1).

**Supplementary Video 5. Visualisation of tight junctions of host IECs from *T. muris* L1 larva in syncytial tunnel in caecaloids.** Video visualising z-stack of confocal IF images of caecaloids infected with whipworm L1 larvae for 24 h. Note tight junctions of infected IECs are conserved. Stained in blue are nuclei of both the epithelia cells and larvae (DAPI), in white is F-actin at cell membrane (phalloidin), in green are mucus vacuoles of goblet cells (UEA/SNA lectins binding mucin glycans) and in red are tight junctions (ZO-1 protein).
